# The attachment rate of *Vibrio anguillarum* strains to microplastics strongly varies with abiotic and biotic factors

**DOI:** 10.64898/2026.01.20.700601

**Authors:** Massot Méril, Wimmer Lukas, Mhamad Aly Moussawi, Hamet Jeanne, Tatjana N. Parac-Vogt, Dailey Lea Ann, Callens Martijn, Bedhomme Stéphanie

## Abstract

Microplastics (MPs), resulting from plastic objects and waste degradation, are increasingly abundant, particularly in marine environments. They exhibit a hydrophobic surface on which biofilms form easily. Metagenomic analyses of these biofilms have revealed that they often contain bacterial species potentially pathogenic to humans or animals. For this reason, MPs are suspected to present a risk for public health by acting as a vector for pathogenic bacteria species. To better understand this hazard, we studied different factors potentially affecting the bacterial attachment rate to MPs. Focusing on the fish pathogen *Vibrio anguillarum,* a collection of 16 strains was assembled and GFP-labelled. Their attachment rates were measured using fluorescence microscopy on three types of MPs (milled polypropylene and polyethylene terephthalate particles and commercially available polystyrene beads). A strong effect of the particle type was found, likely linked to both the chemical composition of the particles and the surface characteristics, with higher attachment rates on rough particles. Our results also revealed a strong intra-specific variation in attachment rate, highlighting the need of testing several strains of the same species to assess attachment rate and related hazards. Finally, it was observed that when a biofilm already formed on the MPs (by field-incubation of the MPs along the Mediterranean French coast), differences in attachment rates between particle types were erased. It was concluded that the attachment rate of *V. anguillarum* depends on a combination of biotic and abiotic factors, which makes risk assessment of MPs as vectors of pathogenic bacteria species a very complex task.

## Introduction

Approximately 9 billion tons of plastic objects have been manufactured since 1950, of which more than half have been produced since the early 2000s [1]. Of all the discarded plastics so far, some 14% have been incinerated and less than 10% recycled [1], whereas the vast majority ends up in various ecosystems. As a result, plastic material can already be found everywhere on the planet [2], [3], [4], [5], and is the largest source of debris produced on the continents and ending up in the oceans [6].

Microplastics (fragments between 1 µm and 5 mm) and nanoplastics (fragments smaller than 1 µm), are considered to be a concerning element of plastic pollution. Studies indicate they can reach virtually all organs of the human body, e.g. the blood [7], the brain [8], the kidneys [8], the placenta [9], and the digestive system [8], [10]. A growing body of literature raises concerns regarding their cytotoxicity [11], [12] and broader health implications [13], [14], [15]. In addition to the negative effects on animal and human health through physical damage and chemical interactions, microplastics can present a risk through the molecules and living organisms that adhere on them. Indeed, when immersed in seawater or freshwater, microplastics are rapidly covered by a layer of proteins and other biomolecules, called the “ecocorona” [16]. This nutrient-rich ecocorona attracts bacteria which can also adhere to the microplastics and form a biofilm. The term ‘plastisphere’ has been coined to describe communities of microbes associated to microplastics [17]. Several studies have detected an enrichment of potential pathogens in biofilms formed on microplastics collected from seawater, compared to the water column [18], [19], [20], [21]. However, many knowledge gaps have been identified and should be investigated to confirm or disprove the assumption of microplastics acting as vectors for pathogens [22].

A first potential issue is that numerous *in situ* studies have detected potential pathogenic bacteria within the plastisphere *via* 16S rRNA gene sequencing [23], [24], [25], [26], [27], [28]. This technique is essential for studying microbial communities, but lacks consistent taxonomic granularity that allows identification at the species level, preventing conclusions about the actual presence of pathogenic species [29]. Even when pathogenic species were detected *via* other techniques, their viability or their virulence was not tested [19], [30], [31]. Moreover, strain identity and diversity within pathogenic species of the plastisphere remain overlooked although there can be a large variability of pathogenicity (ability to cause a disease) and virulence (severity of the disease) within a bacterial species. Intraspecific variability in gene content involved in motility and biofilm production has been described in several bacterial taxa [32], [33], [34]. These functions partially determine the expression of virulence towards a host, as they enable adhesion to its epithelia. Hence, genes involved in the adhesion to surfaces and virulence often belong to the accessory genome and that there is a strong intra-specific variability for the expression of these traits. Although studying intraspecific variability is key to understand bacterial attachment to solid surfaces and biofilm formation, this has received less attention than interspecific variability.

A second potential issue is that the term “microplastics” covers a huge diversity of particles in terms of physical and chemical properties like polymer type, density, shape, additives, surface regularity and chemistry (e.g. hydrophobicity), or fragmentation process [35]. The influence of these properties, alone and in combination, on the potential of microplastics as a vector of pathogenic bacteria has been largely overlooked in previous studies. Furthermore, the majority of microplastics studied to date were collected in marine environments [21], [36], and consequently, these properties were not controlled.

Many reports on enrichment of the plastisphere with potential pathogens concern the bacterial genus *Vibrio*. It contains several species causing severe diseases in humans and marine animals, including the well-known causative agent of cholera, *Vibrio cholerae*. This genus is usually found in the early stages of biofilm formation on microplastic surfaces, where it becomes a minor member of mature biofilms [25], [26], [27]. However, a certain inter-study variability remains.

The objective of this study was to investigate some of the factors driving the attachment of pathogenic bacteria to microplastics. *Vibrio anguillarum* was chosen as the model species, as it is lethal to fish and crustaceans and is responsible for substantial losses in aquaculture. More specifically, four questions were asked: (1) Is there an intraspecific variation of the attachment rate to microplastics in *V. anguillarum* and, if so, can we identify the genomic features behind this variation? (2) Does the attachment rate vary between microplastic types (polymer and shape)? (3) Does the presence of a natural biofilm, formed by in-field incubation, change the attachment rate of *V. anguillarum*? (4) Does the attachment to microplastics increase the resistance of *V. anguillarum* to bleach, a common water sanitizing treatment?

## Material and methods

### 1. Bacterial strain and genomic analyses

#### 1.1. Selection of *V. anguillarum* strains

To explore the intraspecific variability of attachment rates on MPs, 16 *V. anguillarum* strains were obtained from the Belgian Coordinated Collection of Microorganisms (Table S1). These strains were selected to represent a wide geographic range and diverse ecological origins (isolated from infections in various host species or from environmental samples) (Figure 1A).

**Figure 1.**
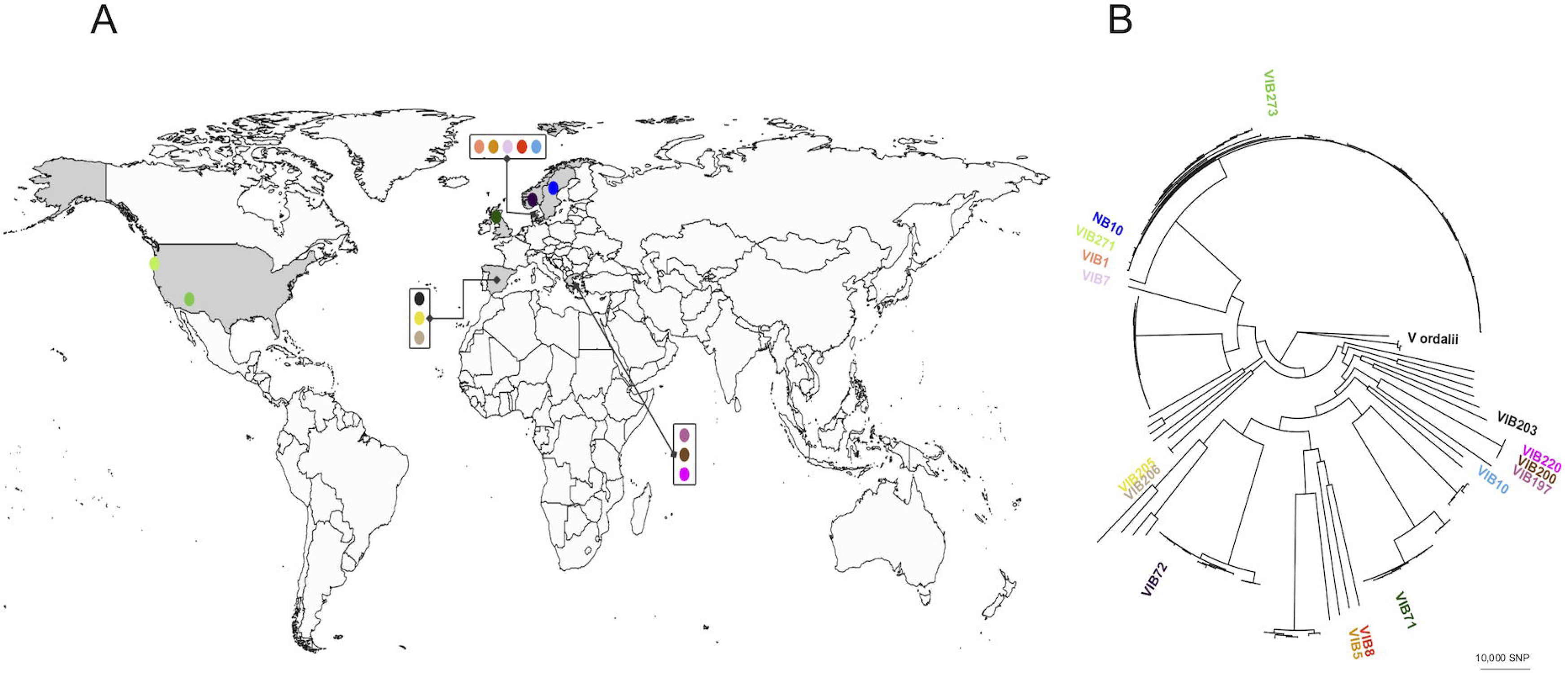
Phylogenetic tree and geographic origin of the tested *V. anguillarum* strains. Panel A represents the geographical origin of each strain. Colors refer to the identity of each strain on the phylogenetic tree. Countries of origin are colored in gray. Panel B represents the maximum likelihood phylogenetic tree of the 156 *V. anguillarum* strains. The tree was rooted on the *V. ordalii* FS-32 strain. It was built using IQ-Tree 2.1.4-beta, and is based on the 2,135 genes that composed the core genome. For clarity purpose, the tree presents only the 16 strains tested in this analysis.

#### 1.2. Fluorescent labelling of *V. anguillarum* strains

The pMRB plasmid [37] containing a green fluorescent protein (GFP) coding gene and a chloramphenicol resistance gene as selection marker was introduced by conjugation in all *V. anguillarum* strains (supplementary material). Briefly, 200 µL of *Escherichia coli* ß3914 (diaminopimelate-auxotrophic donor) containing pMRB-GFP plasmid was mixed with 800 µL of recipient culture, centrifuged and spotted on top of a sterile cellulose acetate filter (0.45 µm pore size). Filters were placed on LBA + NaCl 0.5M supplemented with diaminopimelate (0.3mM). Plates were incubated overnight at 28 °C. Bacterial cells were resuspended in LB + NaCl 0.5M, and *V. anguillarum* transconjugants were selected on LBA + NaCl 0.5M + chloramphenicol (5 µg/ml). For each *V. anguillarum* strain, single colonies were isolated, and their GFP expression was assessed, using the fluorescence microscope Leica DM2000 LED (Leica microsystem, Wetzlar, Germany).

#### 1.3. Genome sequencing and phylogenetic reconstruction

##### Sequencing

We acquired high-quality genome sequences using a hybrid approach that combined long-read data from Oxford Nanopore Technology (ONT, Oxford, United-Kingdom) with high-accuracy short reads from Illumina (San Diego, CA, USA). Genomic DNA was extracted using the QIAGEN Blood and tissue DNA extraction kit (Hilden, Germany) following manufacturer’s instructions on overnight cultures in LB + NaCl 0.5M at 28 °C.

Illumina libraries were prepared using the Nextera XT Flex DNA prep kit, and ONT library with the Native barcoding kit V14 SQK-NBD114, both according to the manufacturer’s instructions (supplementary material).

##### Read processing, assembly and annotation

The quality of raw Illumina reads was controlled using FastQC [38] (Phred score threshold= 30) and adapters and regions with low quality-scores were cleaned with Cutadapt (version 4.0) [39]. ONT reads with quality score lower than 15 were discarded using NanoPack2 [40], [41].

Hybrid *de novo* assembly was done using Hybracter (version 0.11.2) [42] on the filtered reads. Genomes were annotated using PROKKA (version 1.14.6) [43].

##### Phylogenetic reconstruction

All the available *V. anguillarum* genomes were retrieved from the curated collection NCBI RefSeq. Subsequently, we assessed the genome integrity using CheckM (version 1.1.3), focusing on completeness and contamination [44]. The analysis was conducted in December 2024 and yielded 140 genomes in total.

Roary (version 3.13.0) [45] was used to determine the core genome of the 157 strains (140 genomes from RefSeq, our 16 sequenced strains, and the *V. ordalii* FS-238 strain as an outgroup). A phylogeny based on single nucleotide polymorphisms (SNPs) in the core genome was built. All core genes were aligned using MAFFT (version 7.505) with the ‘auto’ option [46]. A core gene nucleotide alignment supermatrix was produced by concatenating all aligned core genes. A SNP distance matrix was obtained from the alignment with SNP-dists (version 0.7.0) [47]. A maximum-likelihood phylogenetic tree was constructed with IQ-Tree 2.1.4-beta [48], and a GTR+F+R9 model of substitution chosen according to Bayesian information criterion obtained with ModelFinder [49].

### 2. Microplastic and non-plastic control materials

Attachment experiments were conducted with irregularly shaped polypropylene (PP) and polyethylene terephthalate (PET) fragments (30-50 μm) produced by milling (supplementary material). The milling procedure described by [50] was modified to obtain smaller PP microplastics. A different previously published method [51] was optimized for the production of PET microplastics. MPs passing through a 50 μm sieve and retained by a 38 μm sieve were collected for subsequent analyses. The particle size distributions of the PP and PET fractions were measured *via* laser diffraction, confirming that the majority of particles fell within the range 30-50 μm (Figure S1). Monodisperse polystyrene (PS) spheres a diameter of 30 µm (Sigma-Aldrich, MO, USA) were utilized as comparator plastic material. The surface roughness of plastic particles was assessed using electron microscopy. Scanning electron microscopy (SEM) micrographs were recorded using a JEOL-6010LV SEM. Before imaging, the samples were coated with a palladium layer using a JEOL JFC-1300 autofine coater under Ar plasma. PS particles exhibited smooth surfaces, whereas PET and PP particles showed rough surfaces with irregular cavities, which were absent in PS beads (Figure 2A).

**Figure 2.**
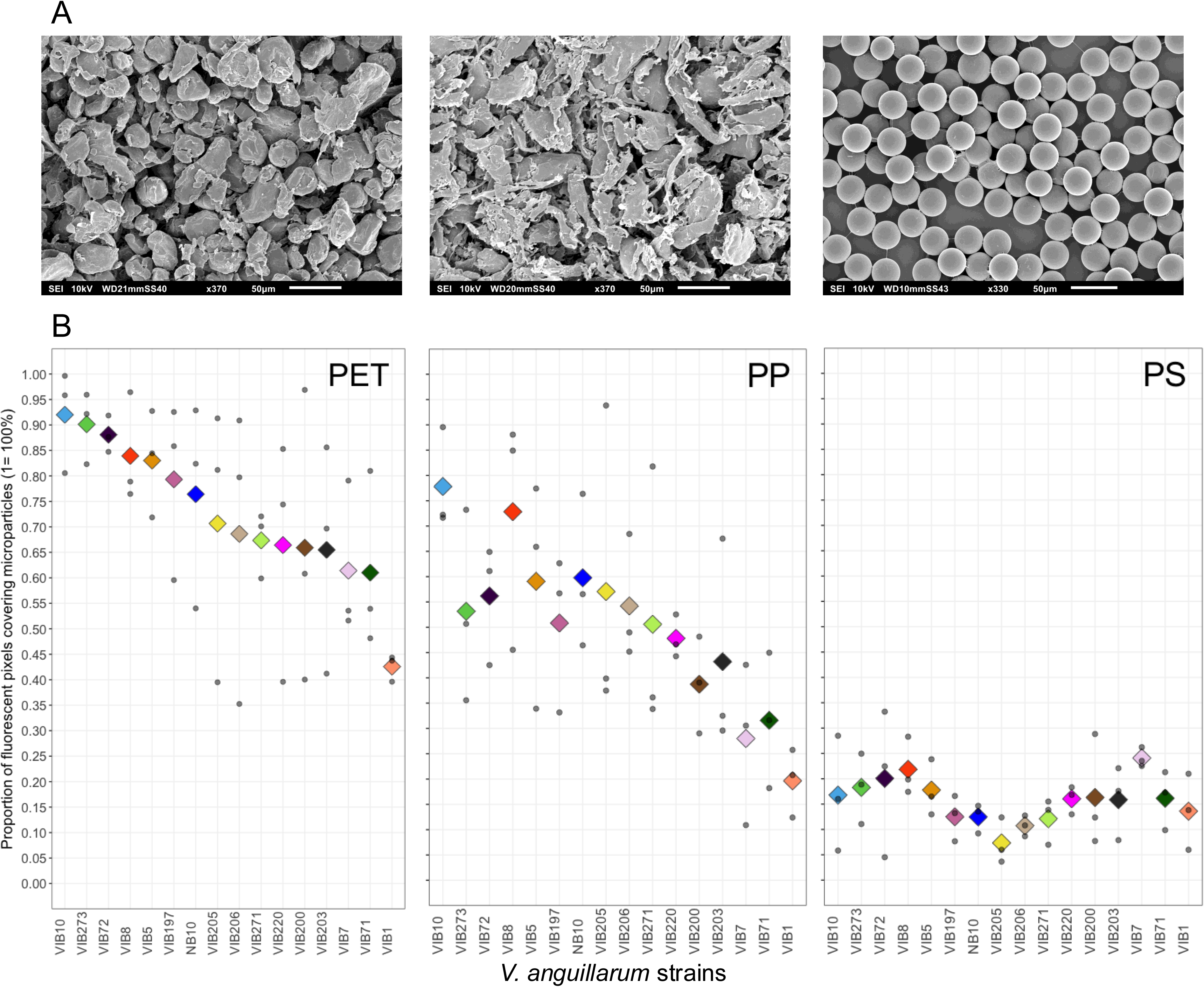
Scanning Electron Microscopy (SEM) images showing microplastic surface morphology, and proportion of fluorescent pixels covering microplastics for the 16 *V. anguillarum* strains tested. Panel A shows SEM images of PET, PP and PS particle surfaces, shown from left to right, respectively. Scale and magnifications are provided at the bottom of each image. Panel B displays the proportion of fluorescent pixels covering the microplastics. Each graph represents attachment data for microplastics from different polymers (from left to right: PET, PP, and PS). For each strain, gray points show the observed data for three replicates, while colored diamonds indicate the mean proportion. The color code corresponds to the strain, and is consistent across all graphs. Strains are organized in descending order based on their mean values for PET.

Glass spheres with a diameter > 108 μm and lignin fragments were used as non-plastic controls (Sigma-Aldrich, MO, USA).

### 3. Experiment 1: *V. anguillarum* attachment on pristine microplastics

#### 3.1. Preparation of bacteria and microplastic mixtures

*V. anguillarum* strains carrying the pMRB-GFP plasmid were grown overnight in LB + NaCl 0.5M supplemented with chloramphenicol (10 μg/mL) at 28 °C, with constant agitation at 200 rpm. Concentrations of the bacteria cultures were determined by flow cytometry absolute counting (supplementary material) and mixtures at a 1:10 ratio of microplastics to bacteria were prepared using microplastic suspensions at an approximate concentration of 5*10^5^ particles per mL. A volume of 1.8 mL of the resulting mixtures were incubated at 30 °C for 24 hours with constant agitation at 200 rpm in 2 mL Eppendorf tubes. Density differences of the polymeric materials compared to the used media caused either flotation (PP) or sedimentation (PET, PS) of the particles. To account for this, the tubes were placed horizontally to maximize the likelihood of interaction between planktonic cells and particles. Each combination of the 16 strains and three plastic polymers (PET, PS and PP) was tested in triplicate.

#### 3.2. Fluorescence microscopy and imaging

After 24h of incubation, 10 μl of each mixture were dropped onto a Malassez cell, which was placed under the fluorescence microscope Leica DM2000 LED after a sedimentation period of 3 minutes. Multiple pictures were captured of 10 fields forming a row or a column of squares (each square representing a volume of 0.01 μl) using a Flexacam C3 (Leica microsystem, Wetzlar, Germany) connected to the microscope. The specific row or column was selected to ensure that between 40% and 60% of the area was covered by MPs, where feasible, and before any fluorescence detection. A brightfield (BF) picture of the MPs and the grid of the Malassez cell were taken, before a green fluorescence filter was employed to detect the GFP signal. Ten pictures (GF pictures) of each field were captured under fluorescence excitation light at intervals of 20 µm in height, thus covering the total distance between the Malassez slide and the coverslip. This series of pictures allowed for comprehensive assessment of all planktonic bacteria and bacteria attached to the MPs within a volume of 0.1 μl. All the pictures were taken at a magnification of x200 and saved as TIFF.

#### 3.3. Image treatment

To achieve high-quality and reproducible image treatment from multiple raw microscopy captures, we developed a pipeline using Fiji (ImageJ version 2.14.0/1.54i) [52]. An overview of the pipeline is depicted in Figure S2. Parameter specifications can be found in Table S2, and green fluorescent images of bacteria attached to microplastics are shown in Figure S3A. Briefly, for each sample, we generated one brightfield megapicture encompassing the 10 fields, and 10 green fluorescent megapictures (one per depth). From that, we segmented them to get a mask for both microparticles and green fluorescent pixels emitted by fluorescent bacteria.

##### Raw image processing and stitching

Raw BF images were converted to an 8-bit grayscale. For PET and PS microparticle images, a bandpass filter was applied to isolate structures in the microparticle size range. For PP images, a variance filter was applied to highlight regions with local variability, and a Gaussian blur was applied to smooth the images. The transformed BF images of the 10 fields were stitched *via* the MIST plugin [53], [54], with pre-stitching for optimal alignment (Table S2, supplementary material). Cropping was performed to center images, normalize pixel count, and remove edge artifacts.

##### Pixel segmentation

Segmentation of MPs employed trained models *via* Fiji’s trainable Weka segmentation plugin [55], [56], a machine learning-based image analysis. The segmentation resulted in binary pictures capturing the position of the MPs on the stitched grayscale transformed BF picture. Each mask produced by its utilization was visually verified by overlaying it onto the original image. Manual corrections were made in cases of local mis-segmentation (Table S2, supplementary material). The GF pictures were split into the three color channels, and only the green channel was retained for further analyses. Fluorescent bacteria were segmented using different methods depending on the polymer type, including machine learning or differential intensity threshold approaches to account for particle autofluorescence (Table S2, supplementary material).

##### Fluorescent pixel classification

Fluorescent pixels were categorized into different groups based on their overlap with microplastic masks. Fluorescent pixels overlapping those of the MPs and present at the same depth (from 200 to 140 μm for PS beads; from 200 to 120 μm for PET) were classified as originating from bacteria attached to the microplastics. For the PP particles, we observed two behaviors in the water column. In some samples, the particles floated at a depth between 120 and 40 μm, while in others, the particles settled at the bottom of the column, between 200 and 120 μm in depth. This difference in behavior between floating and sinking PP was considered during pixel classification. Fluorescent pixels overlapping the microplastics but at shallower depths were treated as noise, as we could not distinguish their origin between a clump of bacteria attached below or a planktonic bacterium at a shallower depth. Fluorescent pixels not overlapping with the MPs were classified as originating from planktonic bacteria, regardless of depth.

Strain attachment to supports was evaluated according to Equation 1, where n_overlap_ was the number of fluorescent pixels overlapping MPs and n_mask_ was the number of pixels in the MP mask).

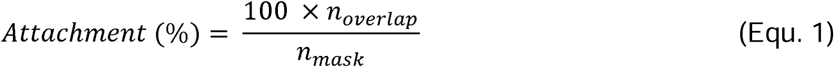

#### 3.4 Exploration of genomic features involved in microplastic attachment

We formulated several hypotheses regarding the potential association of the intraspecific variability in attachment to microplastics with genomic features: attachment efficiency is associated with either (1) intra-species phylogenetic or gene content distances, (2) virulence gene content, or (3) the capacity of bacteria to use plastic polymers as a source of carbon.

First, we performed a phylogenetic reconstruction on the 16 tested strains only, using the same methods as for the complete collection of *V. anguillarum* genomes (see above ‘phylogenetic analyses’) and we computed the phylogenetic variance–covariance matrix obtained from the phylogenetic tree using the vcv.phylo function of the ‘ape’ package [57]. This matrix was later introduced as a random effect in a mixed effect model to account for a phylogenetic influence on attachment to microplastics (see below ‘statistical analyses’ section). Roary was also used to obtain the accessory gene content of each strain. This was later used to test the association between the attachment rate and the presence/absence of genes in the accessory genome (see below ‘statistical analyses’ section).

Second, to determine whether attachment efficiency was associated with virulome content, we constructed a database compiling all experimentally validated genes involved in the virulence of *V. anguillarum*. Briefly, we looked for ‘*Vibrio anguillarum*’ in Pubmed, and read title and abstract of all the articles published between January 1965 and January 2025 to identify relevant studies. Accession numbers corresponding to virulence genes were retrieved from publicly available genomic repositories NCBI [58] and DDBJ [59]. Gene sequences were curated and organized according to their virulence system into a structured database of 308 sequences for further comparative analyses. The dataset used in this study is publicly available at Zenodo (DOI: <u>10.5281/zenodo.18151786</u>). Then, the 16 *V. anguillarum* genome scaffolds were screened for virulence genes using Abricate (version 1.0.1, minimum percent sequence identity: 80%, minimum percent coverage: 90%) [60] against our custom database, as well as the available virulence database ‘vfdb’ (last update: January 2025) [61], [62].

Third, scaffolds were screened for hydrocarbon degradation protein sequences using Proteinortho (version 6.3.4) [63] against the HADEG database (version ‘HADEG_protein_database_231119.faa’) [64], to assess whether certain strains could use plastic polymers as a source of carbon.

### 4. Experiment 2: *V. anguillarum* attachment on field-colonized microplastics

#### 4.1. On-field experimental set-up

##### Incubation sites

Three localities were chosen along the French Western Mediterranean coast: a semi-enclosed lagoon (bassin de Thau, GPS coordinates of the experimental set-up: 43.37897°N, 3.57164°E), a marina (Carnon’s harbor, GPS coordinates: 43.54609°N, 3.97173°E), and a protected area (the Marine Reserve of Cerbères-Banyuls, GPS coordinates: 42°29300 N – 03°08700 E). Each location was selected due to differences in the type and intensity of human activities: the Bassin de Thau is known for its oyster farming and recreational nautical activities, Carnon’s harbor is a marina. Both are highly frequented during summer, whereas access to the marine reserve is restricted all year long.

##### Field-colonized microplastic incubation

We prepared the sampling device as described in [65]. For each incubation, four nylon bags containing 50 mg of PP, PS, PET or glass microparticles were hung with Ty-raps within an oyster cage (Nodus factory, France). The cage was loaded with four 1 kg diving weights and immersed one meter below the surface. The incubation period was four weeks.

At the end of the incubation period, the bags containing MPs were processed in the lab as described in [65]. Briefly, the bags were cut open and the content was resuspended in a sterile artificial seawater (Sea salt, InstantOcean, VA, USA) by aspersion. The obtained suspensions were transferred onto a 20 µm mesh filter to collect MPs and discard the planktonic material. The field-colonized MPs were resuspended in 2 mL of artificial seawater, of which 400 µl were dedicated to *V. anguillarum* attachment rate measurement. Brightfield images of natural biofilms formed around microplastics are shown in Figure S3B.

#### 4.2. Incubation of field-colonized particles with fluorescently labelled *V. anguillarum* strains

##### *V. anguillarum* strain selection

For attachment experiments with microplastics colonized by marine biofilms, strains NB10, VIB8, and VIB205 were chosen based on (1) their genetic distance, each strain being distinct from each other by more than 30,000 SNPs in their core genome (average SNP number among pairs of the 16 strains = 30,443), and (2) their Jaccard distance calculated on their accessory gene content, which was above 0.30 for each pair (average Jaccard distance among pairs of the 16 strains= 0.26).

##### *In vitro* assay and image processing

The number of resuspended microplastics was estimated for each sample by counting the particles under a microscope using a Malassez counting chamber. The incubation protocol and image capturing scheme were similar to the ones described for the first experiment. Briefly, 100 μl of the field-colonized MP suspension was mixed with the required volume of bacteria suspension to achieve a ratio 1:10. The mixture was incubated for 24 hours at 30 °C, with constant agitation at 200 rpm. Each combination of the three *V. anguillarum* strains, three locations, three plastic polymers, and the non-plastic control (PET, PS, PP, and glass) was tested in triplicate. For each combination, we also incubated virgin microparticles in parallel as a paired control under the same conditions.

The same image processing procedure as for the first experiment was employed using our custom Fiji pipelines for PET, PP and PS microparticles. For glass beads, a modified version of the PS microparticle pipeline was developed (Table S2, supplementary material). The outcome variable of this experiment was the difference in the proportion of fluorescent pixels overlapping with particles between field-colonized particles and their paired control.

### 5. Experiment 3: Effect of microplastics on *V. anguillarum* survival to bleach treatment

We tested the effect of biofilm on the survival of *V. anguillarum* to bleach treatment. We used PP and PET microparticles, fragments of lignin (Sigma-Aldrich, MO, USA) as non-plastic control, and VIB205 as focal strain. MPs were mixed with the required volume of bacteria suspension to achieve a ratio 1:10. The mixture was incubated in pre-filtered (0.45 µm) and autoclaved seawater, which was sampled from Carnon’s harbor (GPS coordinates: 43.546231°N, 3.971839°E). Each combination of the focal strain and material (PET, PP, and lignin) was tested in triplicate. We also incubated the focal strain in absence of MPs under the same conditions as a ‘non-biofilm support’ control.

Incubation was conducted in a total volume of 100 mL at 28 °C with constant agitation at 150 rpm to enable *V. anguillarum* cells to attach to MPs. After 5 days of incubation, 2.7 µL of bleach at 37 g L^-1^ were added to each sample (final concentration of 1 mg L^-1^). After 30 minutes of exposure to bleach, each sample was poured on PluriStrainer Mini cell strainer (PluriSelect, Germany) with a mesh size of 20 µm to retain colonized MPs. Biofilms were separated from MPs as described in [66]. According to the track plate method, 10 µL of undiluted resuspension and dilutions from 10^-1^ to 10^-3^ were spotted on TCBS plates and incubated at 28 °C. Survival to bleach treatment was evaluated based on the number of colonies formed after 48 hours.

### 6. Statistical analyses

For the first experiment, attachment was evaluated based on the proportion of fluorescent pixels covering the MP surface. To analyze the effect of polymer type, a generalized linear mixed-effect model (GLMM) was employed. The Beta distribution was used to model the proportion data. Along with polymer type, we also assessed the effect of *V. anguillarum* strains and their interaction. The day of the experiment was included as a random effect to account for variability between replicates.

Model parameters were estimated in a Bayesian framework, using a Markov Chain Monte Carlo (MCMC) algorithm, implemented with the R package brms [67], [68], [69] (supplementary material). The models were compared using the Watanabe-Akaike Information Criterion (WAIC). Among the models with the smallest WAIC values and a difference of less than 4 units, we selected the most parsimonious model. Once the best model was selected, we included the phylogenetic variance–covariance matrix as a ‘random effect’. We tested whether the proportion of variance attributed to the phylogenetic structure significantly contributed to the total variance in the model, and in which proportion.

Next, we tested the association between the attachment rate and the presence/absence of genes in the accessory genome of the 16 strains using Scoary2 (version 0.0.15, Benjamini-Hochberg correction of p-values with a cut-off of 0.05) [70]. To enhance statistical power, we selected candidate genes present in at least 4 strains and no more than 12 strains.

For the second experiment, the difference in the proportions of fluorescent pixels overlapping with particles between field-colonized particles and their paired control was used. Model parameters were estimated in the same Bayesian framework, as described above. The effects of polymer type, location of particle field-colonization, and *V. anguillarum* strains were analyzed using a generalized linear model. The Beta distribution was used to model the data after applying an affine transformation, which mapped the data from the interval [-1, 1] to [0, 1]. The day of the experiment was included as a random effect to account for variability between replicates.

All statistical analyses were conducted using R version 4.4.0 [71].

## Results

### 1. Phylogenetic analyses

We gathered a total of 157 WGS, consisting of 156 *V. anguillarum* strains and one *V. ordalii* strain, which was placed as an outgroup of the phylogenetic tree (Figure 1B). Among these, the genomes (composed of two chromosomes in *V. anguillarum* species) of the sixteen tested strains for MP attachment were assembled *de novo* using hybrid Illumina–ONT sequencing. The average genome size was 4,189,500 bp, with an average GC content of 44.5%. The assemblies consisted of 2 to 10 contigs each, with fourteen genomes resolved into complete circular chromosomes. The hybrid assemblies showed high coverage depth (average >500X), supporting accurate reconstruction of chromosomal structures. The core genome comprised 2,135 genes, with a total of 14,364 genes detected across all genomes, resulting in an average of 3,739 genes per strain. A phylogeny based on single nucleotide polymorphisms (SNPs) in the core genome was built using the *V. anguillarum* genome assemblies from RefSeq, the newly sequenced genomes, and the *V. ordalii* FS-238 strain. The tested strains were distributed throughout the phylogenetic tree, with three clusters of closely related strains observed (< 1,000 SNP): NB10, VIB1, VIB271, and VIB273; VIB197, VIB200, and VIB220; VIB205, and VIB206 (Figure 1B). The average number of SNPs among pairs of tested strains was 30,443.

### 2. Experiment 1: *V. anguillarum* attachment on pristine microplastics

#### Attachment depends on plastic polymer and V. anguillarum strain

Technical issues resulted in missing brightfield and green fluorescence raw images for 10 out of 144 samples. Nevertheless, all affected samples, including those with PS (2 samples), PP (5 samples), and PET (3 samples), were retained in the final analysis (supplementary results, Table S4). After cropping, we compared the size of the reconstructed megapictures, expressed as a number of pixels, to ensure comparability between polymers. No significant difference was detected between polymers (anova, p-value= 0.06, Figure S4).

The mean proportion of fluorescent pixels covering the microplastic surface for PET, PP and PS was 72.6% (± 19.6%), 50.0% (± 21.0%), and 15.7% (± 7.2%), respectively, with a wide distribution for PET and PP (ranging from 35.3% to 99.6% and 10.9% to 93.8%, respectively, Figure 2). For PS, this low proportion combined with very few fluorescent pixels in absolute number was consistent with the almost complete absence of fluorescent bacteria attached to MPs observed by visual inspection under the microscope. Within each polymer, a continuum of attachment was observed among the strains, rather than distinct groups (Figure 2).

Using a generalized linear mixed-effects model, we assessed whether the proportion of fluorescent pixels covering the microplastic surface differed between plastic polymers. The model including polymer type and *V. anguillarum* strain as explanatory variables, with no interaction terms, had the lowest WAIC (Table 1) and was thus considered the best model. Based on the Bayesian model comparison, we found substantial evidence supporting a strong effect of polymers and strains (Table 2, Bayes Factor_model null-model 1_= 0.0006 and BF_model 1-model 2_<10^-5^).

**Table 1.** Model selection for the first experiment. Model in bold correspond to the model selected.

| Model | Covariables | WAIC |
| --- | --- | --- |
| null | null | -133.7 |
| 1 | Plastic polymer | -133.3 |
| **2** | **Plastic polymer**<br>**Strains** | **-162.1** |
| 3 | Plastic polymer<br>Strains<br>Interaction(plastic polymer; strain) | -137.1 |

**Table 2.**
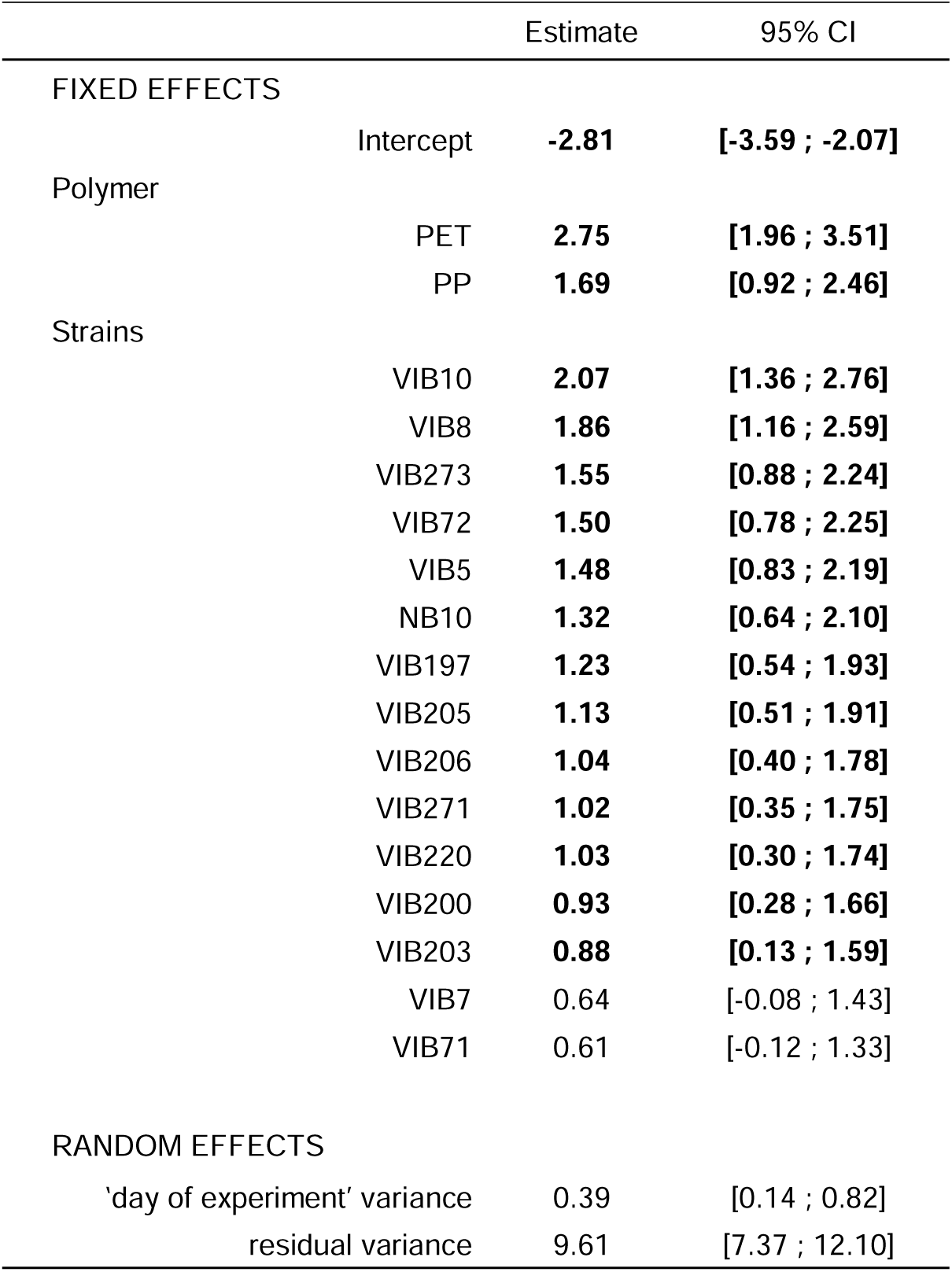
Median and 95% credible intervals of posterior distributions for the estimates of the proportion of fluorescent pixels covering microplastics. Levels used as reference were “PS” and “VIB1” for polymer and V. anguillarum strains, respectively. Estimates in bold correspond to distributions for which the confidence interval does not contain zero.

The absence of significant interaction between strain and polymer in the model retained suggests that strains exhibited similar behavior in their attachment across the three polymers. Despite an important inter-replicate variability, strains with a high attachment phenotype on PET also exhibited a high attachment phenotype on PP. These similarities in ranking between PET and PP were not observed in the presence of PS, where very few attached bacteria were observed.

#### Exploration of genomic features involved in microplastic attachment

To test the hypothesis that differences in attachment efficiency was explained by phylogenetic distances, the variance-covariance matrix derived from the phylogenetic tree of the core genome was introduced as a random effect in the mixed effect model. The proportion of variance attributable to phylogenetic relatedness was 0.03 [0.00; 0.09]_CI95%_, indicating no phylogenetic signal. No significant association was found between any gene of the accessory genome and the attachment gradient to PET or PP.

Among the 307 Vibrio anguillarum-specific gene sequences encompassing 23 virulence systems gathered in our custom database, we detected between 220 and 277 virulence genes (median: 230), with the majority localized on the first chromosome (median number on the first and second chromosome being 167 and 64, respectively). The number of virulence genes was not correlated with their attachment efficiency (rho= −0.13, p= 0.6, Spearman’s correlation test). There was no correlation either when focusing on genes involved in adhesion and quorum sensing (rho= −0.07, p= 0.8, Spearman’s correlation test).

To assess whether the attachment rates of *V. anguillarum* strains to MPs could be linked to the presence of genes encoding plastic degrading enzymes, the annotated genomes were screened for the presence of genes of the HADEG database. The outer membrane lipoprotein Blc (UniProt accession number Q08790) and the protein-tyrosine-phosphatase Wzb (UniProt accession number P0AAB2) were the only sequences detected in 15 and 10 strains, respectively. No association was found for the attachment gradient, since strains without these sequences did not exhibit an outlier phenotype.

### 3. Experiment 2: *V. anguillarum* attachment on field-colonized microplastics

Technical issues resulted in missing brightfield and green fluorescence images for 7 samples out of 216 samples. Nevertheless, all affected samples, including those with PS (2 samples), PET (1 sample), PP (2 samples), and GL (2 samples), were retained in the final analysis (supplementary material). After cropping, there was no significant difference between the size of pictures between naturally-colonized particles and their control, nor between strains and locations. The surface of BF pictures covered by PS beads was significantly lower than for the other substrates (posthoc Tukey test, p-value < 10^-5^ for the three comparisons, Figure S5), as for Thau compared to Carnon incubation sites (posthoc Tukey test, p-value= 0.001).

As a preliminary step, we confirmed that strain attachment to pristine MP particles was consistent with the results of the first experiment (Figure S6). Attachment was highest on PET (81% ± 15.7% on average), intermediate on PP (73.4% ± 20.1%), and lowest on PS (15.9% ± 12.1%). For each of the three strains, the proportion of fluorescent pixels overlapping particles in experiment 2 was in line with that observed in experiment 1, with the largest difference in mean observed for VIB205 on PET (70.7% ± 27.5% on average in experiment 1, 90.6% in experiment 2 ± 8.4%).

For each combination of strain, field location and polymer type, we computed the difference in fluorescent pixel overlap between field-colonized and pristine particles (Figure S7). Positive differences reflected increased attachment of GFP-labelled *V. anguillarum* to microplastics in the presence of naturally formed biofilm, while negative differences reflected decreased attachment.

The difference in fluorescent pixel overlap was negative for PET and PP (PET= −22.3 ± 23.0, PP= −26.5 ± 28.3, Figure 3, Table S5), while positive for PS beads (22.0 ± 27.5). The difference in proportion of fluorescent pixels covering the microplastic surface between naturally colonized microparticles and their paired control was similar among incubation sites (−4.8 ± 22.3, −5.1 ± 36.3, and −7.4 ± 33.9 for Banyuls, Carnon and Thau, respectively, Table S5) and among tested strains (−12.5 ± 31.1, −5.8 ± 33.8, and 0.9 ± 27.7 for NB10, VIB8, and VIB205, respectively, Table S5).

**Figure 3.**
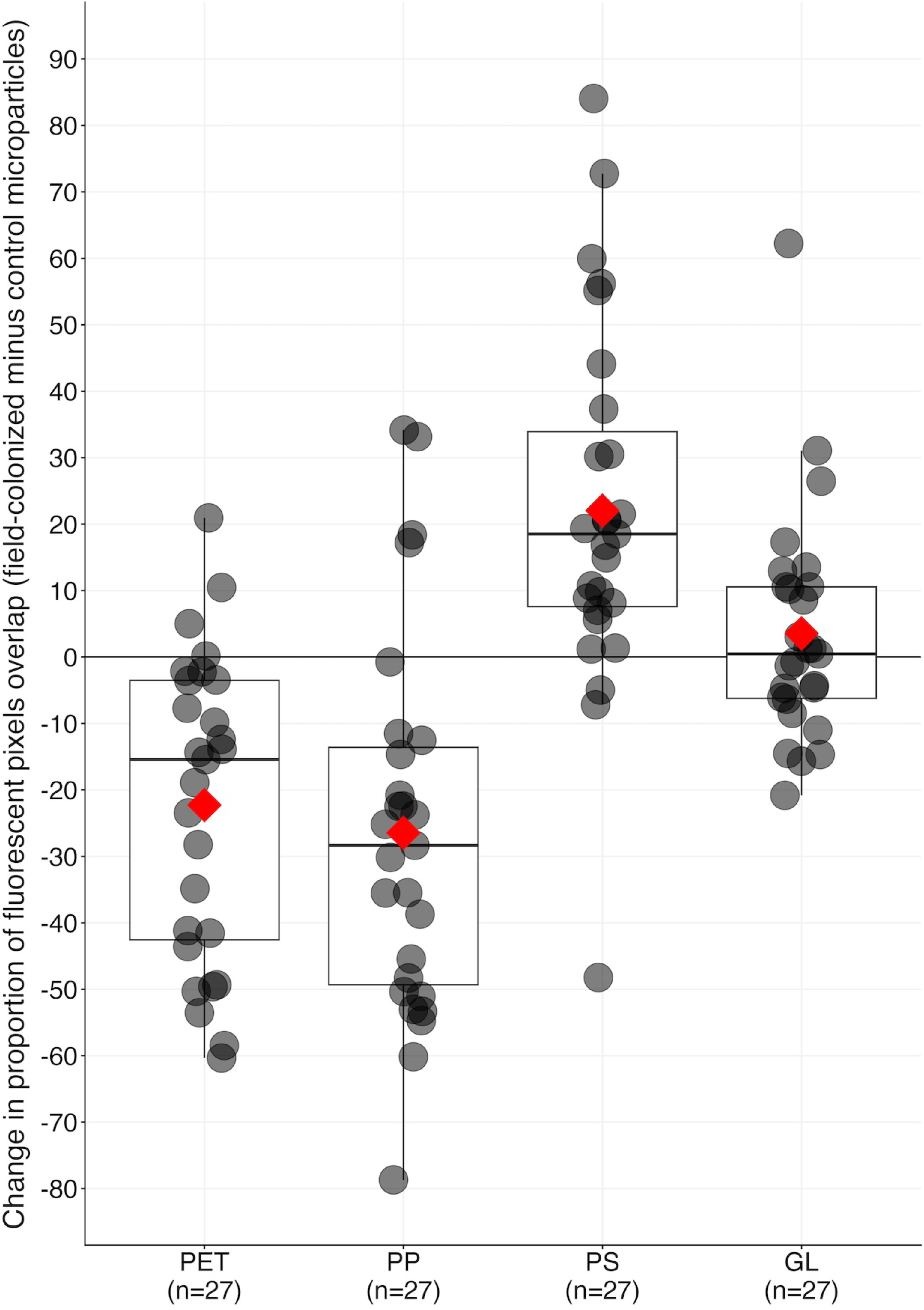
Change in proportion of fluorescent pixels overlap between field-colonized microparticles and control, according to the nature of substrate. For each strain, gray points show the difference between paired samples for each combination of {substrate – location - *V. anguillarum* strain}, while red diamonds indicate the mean difference for each substrate. A difference below zero indicates lower colonization of field-colonized substrate compared to the control by the focal *V. anguillarum* strain.

The model with polymer and *V. anguillarum* strain variables and with an interaction term exhibited the lowest WAIC, while the simpler model that included only the polymer variable differed by less than 4 points (Table 3). Given the principle of parsimony, we selected the model including only the polymer type for further analyses.

**Table 3.** WAIC of models run for the second experiment. Model in bold correspond to the model selected.

| Model | Covariables | No interaction | Interaction |
| --- | --- | --- | --- |
| null | null | -87.5 | - |
| **1** | **Plastic polymer** | **-136.4** | - |
| 2 | Plastic polymer<br>Strain | -138.3 | -135.5 |
| 2 | Plastic polymer<br>Location | -132.4 | -140.0 |
| 3 | Plastic polymer<br>Strains<br>Interaction(plastic polymer; strain) | -135.5 | -118.9 |

In the presence of naturally formed biofilms, *V. anguillarum* showed significantly reduced attachment to PP and PET, whereas attachment was significantly enhanced on PS, compared with the non-plastic support (Table 4, −0.53 [−0.82; −0.26]_CI95%_, −0.62 [-0.89; −0.37]_CI95%_, 0.40 [0.11; 0.68]_CI95%_, for PET, PP and PS, respectively).

**Table 4.**
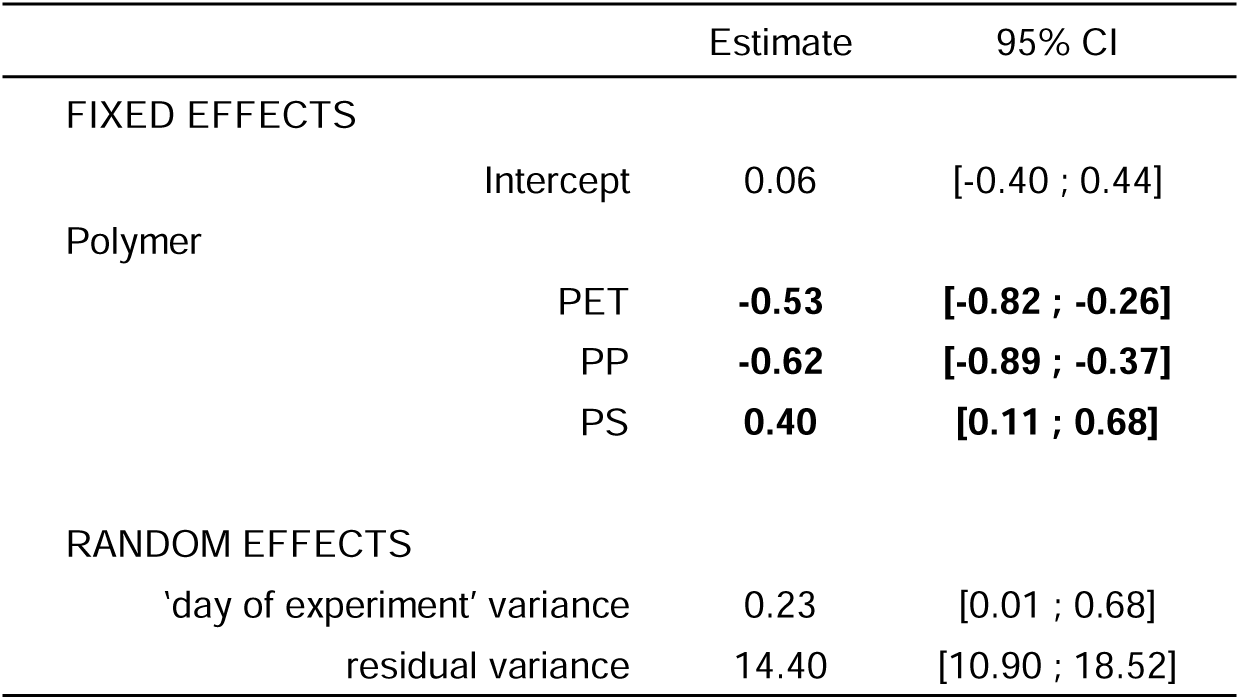
Median and 95% credible intervals of posterior distributions for the estimates of the difference of proportions of fluorescent pixels covering microplastics between field-colonized particles and pristine. Level used as reference was “GL”. Estimates in bold correspond to distributions for which the confidence interval does not contain zero.

### 4. Experiment 3: Effect of microplastics on *V. anguillarum* survival after bleach treatment

We tested the effect of attachment to microplastics on the survival of *V. anguillarum* after 30 minutes of bleach exposure at a concentration of 1 mg L^-1^. As expected, the no-support control conditions showed no *V. anguillarum* growth after bleach treatment, whereas viable colonies formed on TCBS following incubation with PET and lignin in two out of three replicates (Table 5), suggesting that attachment to PET or lignin reduces the efficacy of bleach treatment. Colony counts were higher in lignin samples than in PET: lignin replicates yielded 5.3*10L and 1.5*10^4^ CFU/ml, compared with 900 and 100 CFU/ml in PET replicates. No colony was found from PP.

**Table 5.**
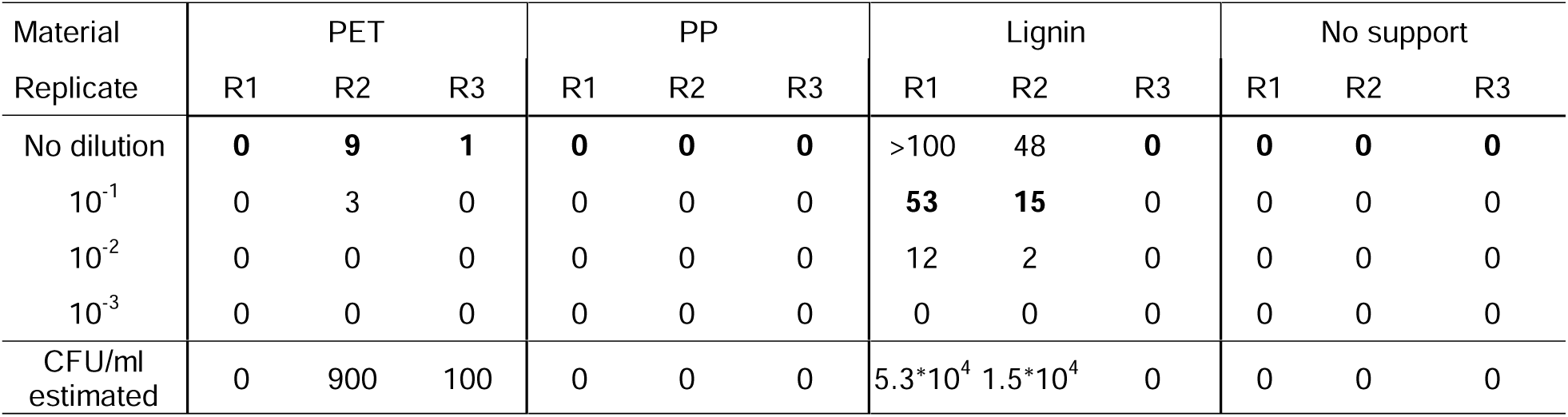
Number of colonies found on TCBS plates after bleach treatment. Numbers in bold were used to estimate the concentration of bacteria after survival.

| Material<br>Replicate | PET |  |  | PP |  |  | Lignin |  |  | No support |  |  |
| --- | --- | --- | --- | --- | --- | --- | --- | --- | --- | --- | --- | --- |
|  | R1 | R2 | R3 | R1 | R2 | R3 | R1 | R2 | R3 | R1 | R2 | R3 |
| No dilution | **0** | **9** | **1** | **0** | **0** | **0** | >100 | 48 | **0** | **0** | **0** | **0** |
| 10<sup>-1</sup> | 0 | 3 | 0 | 0 | 0 | 0 | **53** | **15** | 0 | 0 | 0 | 0 |
| 10<sup>-2</sup> | 0 | 0 | 0 | 0 | 0 | 0 | 12 | 2 | 0 | 0 | 0 | 0 |
| 10<sup>-3</sup> | 0 | 0 | 0 | 0 | 0 | 0 | 0 | 0 | 0 | 0 | 0 | 0 |
| CFU/ml<br>estimated | 0 | 900 | 100 | 0 | 0 | 0 | 5.3*10<sup>4</sup> | 1.5*10<sup>4</sup> | 0 | 0 | 0 | 0 |

## Discussion

The number of scientific studies that show plastics can be major negative disruptors of marine ecosystems keep increasing [15], [72], [73], [74], [75]. One aspect is that plastic debris, and in particular MPs, can be vectors or vehicles for pathogenic bacteria species in marine environments [16]. Here, we explored some of the driving factors for attachment of the fish-pathogen species *Vibrio anguillarum* to MPs. We measured the *V. anguillarum* attachment on PS, PP, and PET microparticles in lab-controlled conditions. This experimental approach was repeated with 16 strains, isolated in both an infection or non-infection context, representing the within-species diversity. We further investigated if the *V. anguillarum* attachment was influenced by a field-acquired biofilm on the MP surface, and whether the effect of attachment to an MP support influences strain survival when exposed to bleach.

Our experimental approach revealed a strong effect of the particle type on the attachment rate of the 16 *V. anguillarum* strains tested (Figure 2B). All three particle types, *i.e.* milled PP MPs, milled PET MPs and commercially manufactured PS beads, were of similar median size (30 µm) but differed in their shape, size range, chemical composition and surface characteristics, e.g. roughness. Furthermore, while the in-house produced PP and PET particles contain no additives, the PS beads might also contain undisclosed additives (PS-beads commercially available but no list of additives was provided). Physically, PS beads have a very smooth surface whereas PP and PET particles obtained by milling have a rough surface with numerous crevices (Figure 2A). The higher attachment rates on PP and PET particles suggest that the physical differences (*i.e.* surface roughness) may explain the “particle type” effect, although a fully crossed design would be necessary to disentangle the effects of roughness and chemical composition on the attachment rate. The positive effect of rugosity on attachment rates has been found in other microplastic and non-microplastic systems. For example, Foulon and collaborators showed that *Vibrio crassostreae* can remain attached for several days on PS microparticles, with longer attachment observed on particles exhibiting greater rugosity [76]. Additionally, Hossain and collaborators showed in a freshwater microcosm that colonization of microplastics by three non-Vibrio species was also correlated to rugosity [77]. The studies analyzing the link between roughness and bacteria attachment rates have been reviewed [78] and there is either a positive relationship between roughness and attachment rate or no relationship. These results outline the importance of the choice of the MP models when designing experiments on bacteria attachment rate to evaluate the risk of pathogen transport by MPs. The use of “realistic” MPs (*i.e.* with similar shape and chemical composition to environmentally aged materials) is key and in this regard, the MPs obtained by milling are more similar to MPs produced by plastic object degradation than PS beads.

We next sought to establish whether there were differences in attachment rates between strains. The strains used in the present study were chosen to be diverse in terms of sampling sites and dates. Furthermore, the obtained whole genome sequences confirmed that they are also spread within the phylogenetic tree of the species, and that they differ greatly in their accessory gene content. Our experimental approach revealed a strong intraspecific variation, with strain-specific attachment rates ranging from 35 to 100% on PET and from 10 to 94% on PP (Figure 2). The intraspecific variability in attachment rates has rarely been tested but measurements performed on other *Vibrio* species [79] revealed important differences between strains involved in biofilm formation on plastic surfaces. The strong intra-specific variability in attachment rates found here highlights that determining the attachment rate of a species is not straightforward and cannot be based on measuring a single “representative” strain. It further means that assessing the risk for a particular MP type as a vehicle of a particular pathogenic species is, in itself, a complex task.

The strong inter-strain variability and the similarity in attachment rates for PET and PP particles suggests that there are genetic determinants of the attachment rate. We tried to identify them by testing associations between the attachment rate measurements and the genomic information contained in the whole genome sequences. However, no significant phylogenetic signal was found and no genomic determinant could be identified. Despite combining a general approach with a more targeted screening of *V. anguillarum* virulence genes from an ad-hoc database (Table S3, 10.5281/zenodo.17525113), we found no evidence that the presence of specific genes, including virulence-associated ones, could account for the differences observed among strains. Our decision to focus on virulence genes was based on the fact that some life-history traits involved in interactions with the environment, particularly adhesion to surfaces, have been shown to depend on virulence factors [80]. This led us to hypothesize that strains with certain virulence genes could have a higher attachment rate, which could, in turn, increase the frequency of virulent or pathogenic strains on MPs. The lack of significant association between genetic composition and attachment rate might be due to insufficient statistical power – given the number of strains used – limited by the tractability of the experimental approach, and image analysis. Another explanation could be non-genetic differences between strains (and even between replicates) in the level of expression of genes involved in quorum sensing, motility or exopolysaccharide synthesis [81], as reported by Foulon *et al*. [76] in a laboratory experiment or Kesy *et al.* [82] in field conditions.

We found that one-month field-colonization in a marine environment altered the attachment rates of three tested *V. anguillarum* strains, such that the effect of the particle type described on pristine particles was mitigated. Attachment rates on PP and PET particles were lower for field-colonized particles than for pristine particles, whereas attachment rates on PS particles were higher for field-colonized particles than for pristine particles (Figure 3). This suggests that the presence of a field-formed biofilm homogenizes surface properties, by masking physical and chemical surface properties. Such patterns could also or additionally evolve due to ecological interactions including both attraction and repulsion cues from the microorganisms of the field-formed biofilms [83]. The dynamics of the plastisphere assembly may largely depend on a combination of stochastic encounters between microplastics and marine bacteria, differences in composition of the ecocorona between polymers [84], differential opportunistic colonization strategies mediated by environmental conditions [79], as well as differential capacities to adhere to surfaces of varying rugosity. The abundance of early colonizers could therefore be determined by the order of attachment resulting from the sum of these forces, whereby species that settle first may gain a competitive advantage or facilitate the subsequent attraction of other species. Such variability could explain long-term changes that are often observed in plastisphere composition dynamics [85], [86].

Biofilm formation is known to increase the phenotypic resistance of bacteria to biocides such as antimicrobials [87] and disinfectants [88]. Since MPs provide surfaces for biofilm development, their presence in aquatic environments may reduce the effectiveness of sanitization measures. Among these, we selected bleach treatment as a widely employed strategy for sewage disinfection and water potabilization. We examined whether MP pollution could reduce its efficacy by facilitating the persistence of pathogenic bacteria. We found that PET microparticles enabled survival of *V. anguillarum* during 10 minutes bleach treatment, as did lignin particles. Further validation of these results will require repeating the experiment with a larger number of replicates, as well as using older and more naturally developed biofilms. Despite a large number of zero values resulting from bacterial mortality caused by the treatment, they stress the need to explore the potential role of MPs as shelter during bleach treatment. To our knowledge, this is the first study that aims to quantify the additional risk of MPs over bacterial survival to bleach treatment.

In conclusion, our experimental approach revealed that the adherence of *V. anguillarum* to MPs, and as a consequence the risk for MPs being a vector of pathogenic *V. anguillarum* strains depends on many factors, including physico-chemical particle surface properties, the presence of other microorganisms, and also the identity of the strain. This combination of factors, and of potential others not studied here, makes the evaluation of the risk a very complex task.

Concerns have also been raised regarding the enrichment of antibiotic-resistant bacteria in biofilms associated to microplastics compared to free-living bacteria [89]. However, studies on non-plastic suspended matter in freshwaters have likewise reported an enrichment of mobile genetic elements linked to antibiotic resistance [90]. This further underscores the importance of elucidating the colonization dynamics of pathogenic species on these surfaces. Pathogenic species that ride plastic litter may disrupt fragile ecosystems, already pressured by overfishing and other types of pollutions. While it is relatively rare for a non-native species to successfully survive in a new environment, rafting on ubiquitous plastic debris could potentially increase the frequency of such events.

## Acknowledgments

We thank Frédérique Le Roux, Delphine Destoumieux-Garzon, Guillaume Charrière, and François Delavat for providing conjugation protocol and plasmids.

We are grateful to the ImpTox project’s members, in particular Andreja Rajkovic and Elsa Gadoin for useful discussions. We also thank Tim Janicke for useful discussions, and Vahiniaina Andriamanga and Marie-Pierre Dubois for their assistance in ONT sequencing.

We also thank MARBEC unit (Univ Montpellier, CNRS, IFREMER, IRD, Montpellier, France) and Ifremer institute (Unité observation et écologie de la restauration des écosystèmes littoraux, Laboratoire Environnement Ressources Occitanie, Sète, France) for giving us authorization to install the incubation device on Ifremer’s experimental oyster farm in Bassin de Thau, in particular Delphine Serais, Hervé Violette. We thank the scientific council of the marine reserve of Cerbères-Banyuls (Réserve Naturelle Marine de Cerbère-Banyuls, Département des Pyrénées-Orientales, France), for providing authorizations to incubate the device on site, in particular Virginie Hartmann. We also thank the Banyuls Observation Sea Service (BOSS FR3724, Observatoire Océanologique de Banyuls, Banyuls-sur-mer, France), in particular Laurent Zudaire. We thank Morgann Michel and the harbor’s office for providing authorizations to incubate the device in Carnon’s harbor as well.

All authors were supported by Horizon 2020 grant (grant number: 965173). We report no conflict of interest. IA tools were used to help improve English. Genome assemblies have been deposited in the European Nucleotide Archive (ENA) under BioSample accession numbers ERS28308090 to ERS28308105, with corresponding assembly accession numbers ERZ28769549 to ERZ28769564. All scripts used for the analyses have been deposited on Codeberg and are openly accessible at the following link: https://codeberg.org/merimas/ImpTox_project_Vibrio_anguillarum_attachment_to_microplastics The virulence gene database and the fasta file used as reference database are openly available at Zenodo (DOI: <u>10.5281/zenodo.17525113</u>, DOI: <u>10.5281/zenodo.18151786</u>).

## Declaration of generative AI and AI-assisted technologies in the manuscript preparation process

During the preparation of this work the authors used ChatGPT in order to improve language. After using this tool, the authors reviewed and edited the content as needed and take full responsibility for the content of the published article.

## CRediT author statement

Méril Massot: Data curation, Formal analyses, Investigation, Software, Validation, Writing-Original draft preparation

Lukas Wimmer: Data curation, Investigation, Resources, Validation, Visualization, Writing – review & editing

Mhamad Aly Moussawi: Resources, Visualization, Writing – review & editing Jeanne Hamet: Data curation, Resources, Validation, Writing – review & editing Tatjana N. Parac-Vogt: Funding acquisition, Methodology, Project administration, Supervision, Writing – review & editing

Lea-Ann Dailey: Funding acquisition, Methodology, Project administration, Supervision, Writing – review & editing

Martijn Callens: Conceptualization, Funding acquisition, Methodology, Resources, Writing – review & editing

Stéphanie Bedhomme: Conceptualization, Funding acquisition, Methodology, Project administration, Resources, Supervision, Writing – original draft, Writing – review & editing,

---

**Table S1.** Information on the origin of *V. anguillarum* strains used for experiments. Information was retrieved from the Belgian Coordinated Collection of Microorganisms catalogue.

| Strain | LMG name | Serotype | Sample origin | Country of origin | Year indicated in BCCM |
| --- | --- | --- | --- | --- | --- |
| NB10 | LMG 32357 | O1 | rainbow trout (*Oncorhynchus mykiss*) | Sweden | 1990 |
| VIB 1 | LMG 10861 | O1 | rainbow trout (*Oncorhynchus mykiss*) | Denmark | 1990 |
| VIB 10 | LMG 10870 | O10 | cod (*Gadus morhua*) | Denmark | 1990 |
| VIB 197 | LMG 13224 | VaNT7 | rotifer | Greece | 1991 |
| VIB 5 | LMG 10865 | O5 | cod (*Gadus morhua*) | Denmark | 1990 |
| VIB 7 | LMG 10867 | O7 | eel (*Anguilla anguilla*) | Denmark | 1990 |
| VIB 71 | LMG 4411 | O2a | Young sea trout (*Salmo trutta*) | Scotland | 1960 |
| VIB 72 | LMG 4437 | O2a | cod (*Gadus morhua*), ulcerous lesion | Norway | 1956 |
| VIB 8 | LMG 10868 | O8 | cod (*Gadus morhua*) | Denmark | 1990 |
| VIB200 | LMG 13227 | VaNT7 | sea bream (*Sparus aurata*), larvae | Greece | 1991 |
| VIB203 | LMG 13230 | O6 | sea bream (*Sparus aurata*), larvae | Spain | 1991 |
| VIB205 | LMG 13232 | O6 | water | Spain | 1991 |
| VIB206 | LMG 13233 | O6 | rotifer | Spain | 1990 |
| VIB220 | LMG 13247 | unknown | water | Greece | 1991 |
| VIB271 | LMG 13583 | O1 | chinook salmon (*Oncorhynchus tshawytscha*) | Oregon, United States | 1974 |
| VIB273 | LMG 13585 | O1 | rainbow trout (*Oncorhynchus mykiss*) | Arizona, United States | 1968 |

**Table S2.** Fiji’s operation details. Each step is presented with specific parameters for each type of microparticle. Different strategies were applied due to heterogeneity in microparticle density and opacity.

| Analysis step | PET | PP | PS | GL |
| --- | --- | --- | --- | --- |
| 1. Raw image processing - BF | 1. converted to 8-bit grayscale<br>2. bandpass filter to filter out structures larger than 100 pixels and smaller than 20 pixels, with a tolerance of 5 | 1. converted to 8-bit grayscale two copies of the image<br>2. variance filter with a radius of 1 pixel applied to copy 1<br>3. Gaussian blur with a sigma value of 5 applied to copy 2<br>4. Copies 1 & 2 merged | 1. converted to 8-bit grayscale<br>2. bandpass filter to filter out structures larger than 100 pixels and smaller than 20 pixels, with a tolerance of 5 | 1. converted to 8-bit grayscale<br>2. bandpass filter to filter out structures larger than 100 pixels and smaller than 20 pixels, with a tolerance of 5 |
| 1. Raw image processing - GF | 1. split into the three color channels<br>2. green channel retained |  |  |  |
| 2. Stitching | 1. Each raw image was copied and converted to an 8-bit grayscale<br>2. Variance filter applied with a radius of 3 pixels to enhance regions with high pixel variability.<br>3. Threshold applied to refine the segmentation: pixels with intensity values between 10 and 255 considered as foreground, while those below 10 were designated as background.<br>4. Images converted to binary masks. The background was set to black.<br>5. MIST plug-in and generation of a text file containing the relative coordinates of images was generated and saved for later assembly<br>6. Stitching of the 10 images transformed for later segmentation using the MIST plug-in feeded with the text file containing the relative coordinates of images |  |  |  |
| 3. Cropping | Horizontal megapictures: rectangular region for cropping starting at position {x=0, y= 110}. This rectangle uses the full image width, and adjusted height to 890 pixels<br>Vertical megapictures: rectangular region for cropping starting at position {x=300, y= 0}. This rectangle uses the full image height, and adjusted width to 1,200 pixels |  |  |  |
| 4. Segmentation - BF | 1. Trainable Weka segmentation classifiers n = 3<br>2. Series of Dilate and Erode to fill masks | 1. Trainable Weka segmentation classifier n = 1<br>2. Series of Dilate and Erode to fill masks | 1. Trainable Weka segmentation classifier n = 1<br>2. Series of Dilate and Erode to fill masks | 1. Trainable Weka segmentation classifier n = 2<br>2. Series of Dilate and Erode to fill masks |
| 4. Segmentation - GF | Vibrio cells attached to microplastic: Setting intensity threshold to 90<br>Planktonic Vibrio cells: removing microparticle areas with brightfield mask, then setting intensity threshold to 82 | Setting intensity threshold to 82 | Trainable Weka segmentation classifier n = 1 | Trainable Weka segmentation classifier n = 1 |
| 5. Image operations | Apply AND operation between microparticle mask and fluorescent pixel masks originating from 120-200 $\mu\text{m}$ depths' pictures | Apply AND operation between microparticle mask and fluorescent pixel masks originating from:<br>-120-200 $\mu\text{m}$ depths' pictures (sinking particles)<br>- 40-120 $\mu\text{m}$ depths' pictures (floating particles) | Apply AND operation between microparticle mask and fluorescent pixel masks originating from 140-200 $\mu\text{m}$ depths' pictures | Apply AND operation between microparticle mask and fluorescent pixel masks originating from 100-200 $\mu\text{m}$ depths' pictures |

**Table S3. Database accession identifiers of *V. anguillarum* virulence genes.** Database compiling experimentally validated genes involved in the virulence of *V. anguillarum*, updated as of January 2025.

**Table S4. Summary of samples that were not meeting requirements after image pipeline analysis.** Each sheet corresponds to a separate experiment and batches of samples are separated by an empty line for clarity purposes. Deviations from the pipeline description are indicated for each sample.

**Table S5.**
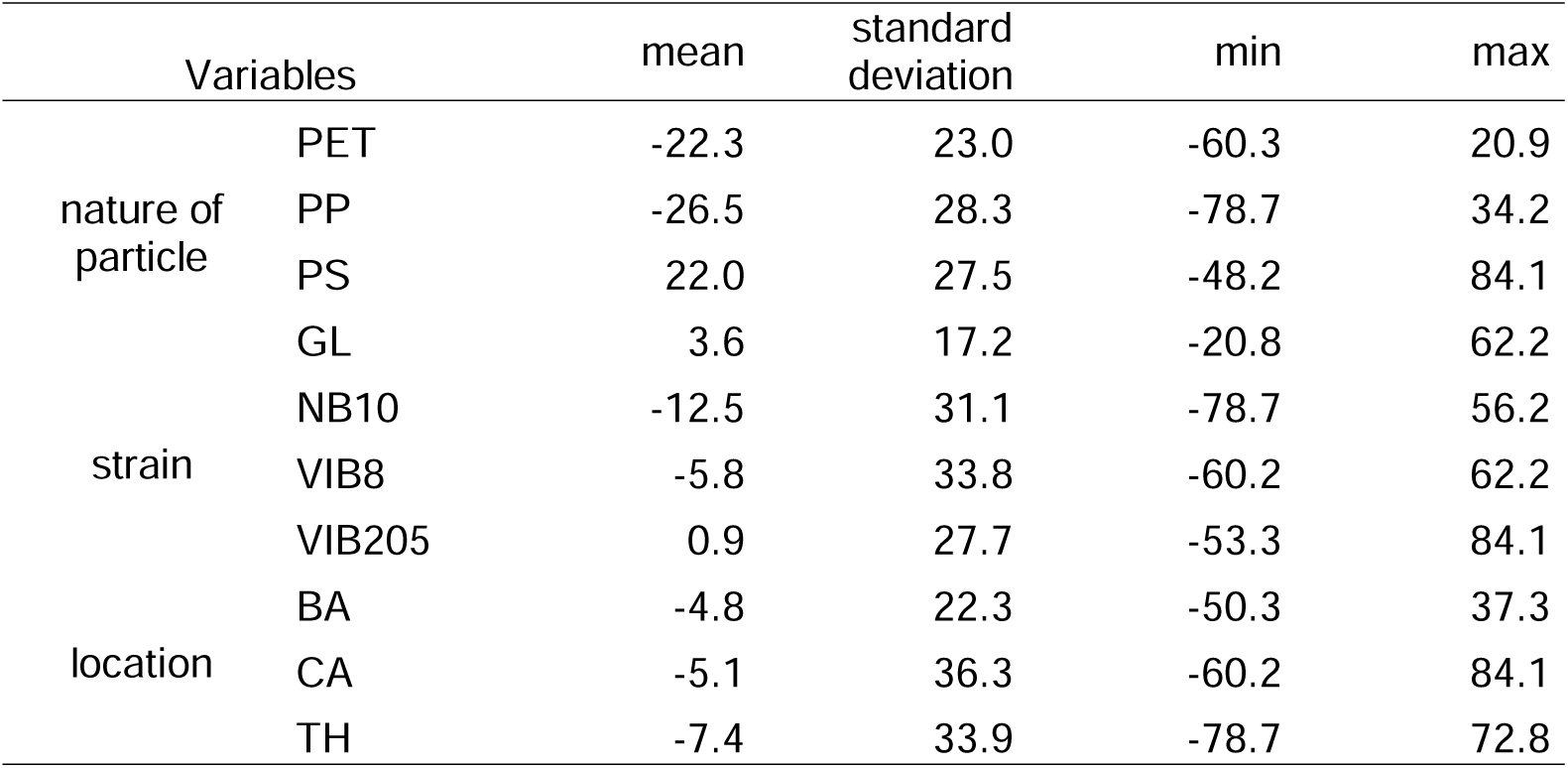
Descriptive statistics of fluorescent pixel overlap on microparticles: comparison between field-colonized and control particles, by nature of particles, *V. anguillarum* strain, and origin of the field-colonized particles. Negative values indicate lower colonization by the focal *V. anguillarum* strain of field-colonized substrate compared to the control.

| Variables |  | mean | standard deviation | min | max |
| --- | --- | --- | --- | --- | --- |
| nature of particle | PET | -22.3 | 23.0 | -60.3 | 20.9 |
|  | PP | -26.5 | 28.3 | -78.7 | 34.2 |
|  | PS | 22.0 | 27.5 | -48.2 | 84.1 |
|  | GL | 3.6 | 17.2 | -20.8 | 62.2 |
| strain | NB10 | -12.5 | 31.1 | -78.7 | 56.2 |
|  | VIB8 | -5.8 | 33.8 | -60.2 | 62.2 |
|  | VIB205 | 0.9 | 27.7 | -53.3 | 84.1 |
| location | BA | -4.8 | 22.3 | -50.3 | 37.3 |
|  | CA | -5.1 | 36.3 | -60.2 | 84.1 |
|  | TH | -7.4 | 33.9 | -78.7 | 72.8 |

**Figure S1. Particle size distribution curve for PET and PP microparticles used in this work.** Individual lines represent the particle size distribution of one batch.

**Figure S2. Pipeline depicting megapicture reconstruction and pixel classification.** Panel A represents image capture protocol of MP/Vibrio cells suspension. Ten microliters were placed on a Malassez cell and a brightfield image of the field was taken to capture microparticle positions at a magnification of x200. A green fluorescence filter was applied to capture GFP signal, and 10 pictures of the field were taken at different heights in 20 µm intervals, encompassing the total volume of the mixture between the slide and the coverslip. Multiple pictures were captured of 10 fields, representing a grand volume of 0.01 μl for each sample. Panels B, C, D, and E represent the different steps using Fiji to generate one brightfield megapicture encompassing the 10 fields, and 10 green fluorescent megapictures (one per depth). From that, we segmented them to get a mask for both microparticles and green fluorescent pixels emitted by fluorescent bacteria. Panel F represents step during which all masks were manually checked and corrected in cases of local mis-segmentation. Panel G represents fluorescent pixel classification, based on their position compared to microplastic masks: (1) those overlapping and at the same depth were classified as emitted by attached bacteria, (2) those not overlapping were classified as planktonic bacteria, (3) those overlapping but at shallower and deeper depths were considered noise. Attachment was evaluated based on the proportion of fluorescent pixels covering the microplastic area.

**Figure S3. Biofilm colonization of microplastic particles observed by light microscopy.** Horizontal bars represent a scale of 10 µm in all images. Panel A shows green fluorescent microscopy images of PS and PET particles after 24 hours if incubation with GFP-labelled *V. anguillarum* in experiment 1. Panel B shows brightfield microscopy images of microplastics after one month incubation along the Mediterranean coast, in experiment 2. Arrows indicate the boundaries of biofilms associated with particles. ‘MP’= microplastic particle

**Figure S4. Number of pixels analyzed in experiment 1 in reconstructed megapictures based on the polymer (*V. anguillarum* attachment on pristine microplastics).** Left panel shows the size of reconstructed megapictures as a function of the polymer, expressed as a number of pixels. The middle panel shows the surface of microparticles covering pictures, while the right panel shows the total number of fluorescent pixels as a function of polymer.

**Figure S5. Number of pixels analyzed in experiment 2 in reconstructed megapictures, based on the polymer (*V. anguillarum* attachment on field-colonized microplastics).** Left panel shows the size of reconstructed megapictures as a function of the polymer, expressed as a number of pixels. The middle panel shows the surface of microparticles covering pictures, while the right panel shows the total number of fluorescent pixels as a function of polymer.

**Figure S6. Comparisons of proportions of fluorescent pixels covering pristine microplastics between experiments.** Each graph represents attachment data for microplastics from different polymers (from left to right: PET, PP, and PS) and strains tested in both experiments (from top to bottom: VIB8, NB10, and VIB205). For each strain, gray points show the observed data, while colored diamonds indicate the mean proportion. Nine measurements per strain and polymer were obtained in experiment 1, and 27 measurements in experiment 2.

**Figure S7. Change in proportion of fluorescent pixels overlap between field-colonized microparticles and control, according to the tested *V. anguillarum* strain, nature of substrate (vertical panels), and origin of the field-colonized particles (horizontal panels).** A difference below zero indicates lower colonization by the focal *V. anguillarum* strain of field-colonized substrate compared to the control. For each strain, gray points show the observed data for three replicates, while colored diamonds indicate the mean proportion. The color code corresponds to the strain, and is consistent across all graphs.

