## Supplemental Material for "The attachment rate of *Vibrio anguillarum* strains to microplastics strongly varies with abiotic and biotic factors"

Supplementary Material and Methods

### 1. Bacterial strain and genomic analyses

##### 1.1. Selection of *V. anguillarum* strains

To explore the intraspecific variability of attachment rates on microplastics (MP), 16 *V. anguillarum* strains were obtained from the Belgian Coordinated Collection of Microorganisms (BCCM). The strains were received in April and November 2021 as freeze-dried material. They were revived in LB + NaCl 0.5M at +28°C for 24 hours, and glycerol stocks were frozen at -70°C for future experiments.

##### 1.2. Fluorescent labelling of *V. anguillarum* strains

The pMRB plasmid [1] containing a green fluorescent protein (GFP) coding gene and a chloramphenicol resistance gene as selection marker was introduced by conjugation in all *V. anguillarum* strains.

###### Bacterial strains and growth conditions

The diaminopimelate-auxotrophic *Escherichia coli* strain ß3914 containing pMRB-GFP plasmid was used as the donor strain. It was cultured in Luria-Bertani (LB) broth and on LB-agar (LBA) supplemented with chloramphenicol (10-25 μg/ml), and diaminopimelate (0.3 mM) at 37 °C with constant agitation at 200 rpm overnight. Recipient *V. anguillarum* were grown in LB broth or on LBA with NaCl 0.5M, at 28 °C with constant agitation at 200 rpm overnight.

###### Bimating conjugation procedure

A 200 µL aliquot of donor culture was centrifuged at 6,500 rpm for 3 minutes, and the supernatant was removed. An 800 µL aliquot of recipient culture was added. The mix was centrifuged, and supernatant was discarded. The cell suspension was gently put on top of a sterile cellulose acetate filter (0.45 µm pore size), on a LBA + NaCl 0.5M supplemented with diaminopimelate (0.3mM). Plates were incubated overnight at 28 °C.

Bacterial cells were resuspended in LB + NaCl 0.5M, and selection for chloramphenicol resistant recipients *V. anguillarum* was done by plating the suspension on LBA + NaCl 0.5M + chloramphenicol 5 µg/ml. For each *V. anguillarum* strain, single colonies were isolated, their GFP expression was assessed, using a fluorescence microscope Leica DM2000 LED (Leica microsystem, Wetzlar, Germany).

##### 1.3. Bacteria quantification using flow cytometry

Serial dilutions from 10^-1^ to 10^-4^ of Green Fluorescent Protein (GFP)-labeled bacteria were analyzed in a Guava® easyCyte 11HT (EMD Millipore Corporation, California, USA) to determine the bacterial concentration per milliliter in the 10^-1^ dilution. A combination of the FSC/SSC gate and of the Green-B channel gate (FITC – excitation 488nm – emission 525/30nm) was set to count the number of fluorescent bacterial cells.

##### 1.4. Genome sequencing and phylogenetic reconstruction

###### Sequencing

We acquired high-quality genome sequences using a hybrid approach that combined long-read data from Oxford Nanopore Technology (Oxford, United-Kingdom) with high-accuracy short reads from Illumina (San Diego, CA, USA). Genomic DNA was extracted using the QIAGEN Blood and tissue DNA extraction kit (Hilden, Germany) following manufacturer’s instructions on overnight cultures in LB + NaCl 0.5M at 28 °C.

We prepared the Illumina libraries using the Nextera XT Flex Illumina DNA prep kit following the manufacturer’s instructions. These libraries were sequenced at Genoscreen (Lille, France) on a Novaseq sequencer (2x150 pb).

We prepared the ONT library using the ONT Native barcoding kit V14 SQK-NBD114 following the manufacturer’s instructions. The prepared libraries were loaded onto an R10.4.1 flow cell and sequenced on a MinION Mk1B device. During the sequencing run, the following quality control metrics were monitored: a minimum Q score of 10, a pore scan frequency of 1.5 hours. Basecalling was performed in real-time by Dorado (version 0.7.4) using the super-accurate basecalling model v4.3.0 at 400 bps.

### 2.  Microplastic and non-plastic control materials

2. 1. Preparation of polypropylene microplastics

Food-grade PP and PET granules were kindly provided by PlasticsEurope. The milling procedure described by [2] was modified to obtain smaller PP microplastics. Eight PP granules (~200 mg) per chamber of a silicone mold (10x10x10 mm) were molded into cuboids (10x10x3 mm) at 180°C for 1 hour. The obtained cuboids were then frozen overnight (-70°C) to increase brittleness. The frozen material (20 g) was then milled in ethanol (96%, v/v, 40 mL) using a knife mill (GM200, Retsch GmbH, Haan, Germany) with cryogenic stainless-steel accessories (Cat. No.: 22.354.0005). Milling with dry ice and subsequent wet sieving of the ethanolic suspension was conducted as described by [2] but the 63 µm sieve was exchanged with a 50 µm sieve (Retsch GmbH, Haan, Germany) resulting in the following sieve stack (from bottom to top): collection pan, 38 µm, 50 µm, 100 µm, 180 µm, 250 µm, and lid. Microplastics retained by the 38 µm sieve (passing the 50 µm sieve) were collected with ethanol and filtered through a 10 µm nylon mesh filter (NY1004700, Merck Millipore KGaA, Billerica, USA) to remove excess ethanol and fine particles. Finally, the microplastics on the filter were dried under reduced pressure using a rotary evaporator (Heidolph Instruments GmbH & Co. KG, Schwabach, Germany) and stored in sealed glass vials (20 mL).

2. 2. Preparation of polyethylene terephthalate microplastics

The higher glass transition temperature of PET allowed milling *via* an ultra-centrifugal mill (ZM 200, Retsch GmbH, Haan, Germany). A similar method for producing PS nanoplastics has been published by [3] and has been optimized for the production of PET microplastics. PET granules (20 g) were pre-cooled in liquid nitrogen for 30 minutes and subsequently mixed with dry ice. The mill was operated at 18,000 rpm (17,963 g) utilizing an 80 µm ring sieve with trapezoid holes and a cyclone with filter bag. The granules in dry ice were transferred spoon-wise into the mill and the milled particles were subsequently collected in a glass bottle by rinsing all parts of the mill with ethanol. The ethanolic suspension was sieved, filtered, dried, and stored as described above.

2. 3. Characterization of microplastics

The particle size distribution (PSD) of the PP and PET materials were measured *via* laser diffraction (Mastersizer 3000, Malvern Panalytical, Malvern, United Kingdom). Briefly, 10-20 mg of powder was mixed with 1 mL of a Tween^®^80 solution (5%, w/w) and bath sonicated and vortexed for 1 minute each. The suspension was added in 100 µL increments to the measurement cell (HydroSV) filled with 6 mL of a Tween^®^80 solution (0.3%, v/v) until a laser obscuration (optical concentration) of 5-10% was achieved. Instrument settings included: Particle shape: “non-spherical”, Mie-Theory was applied (refractive index of PP: 1.490 and PET: 1.636, absorption of PP and PET: 0.01, refractive index of water: 1.33), and stirring speed: 1,200 rpm. The background was measured with 6 mL of distilled water containing 0.3% Tween^®^80. Ten measurements were conducted at room temperature and subsequently averaged.

### 3.  Image treatment

3. 1. Raw image processing

To achieve high-quality and reproducible image treatment from multiple raw microscopy captures, we developed a pipeline using Fiji (Table S2, Figure S1, ImageJ version 2.14.0/1.54i) [4]. Raw BrightField (BF) images were converted to 8-bit grayscale to standardize intensity values.

For PET and PS microparticle images, a bandpass filter was then applied with parameters set to filter out structures larger than 100 pixels and smaller than 20 pixels, with a tolerance of 5. The goal was to isolate structures in the microplastic size range while reducing background noise.

For PP microparticle images, a variance filter with a radius of 1 pixel was applied to highlight regions with local variability. Then, a Gaussian blur with a sigma value of 5 was applied to smooth the images. Subsequently, the results of these two image processing methods were merged, which improved the segmentation of microparticles compared to using either method alone.

3. 2. Getting stitching coordinates

Stitching involved associating the BF images of the 10 fields using predetermined coordinates, with pre-stitching for optimal alignment. Briefly, each raw image was converted to an 8-bit grayscale format to standardize the intensity values across all images. A variance filter was applied with a radius of 3 pixels to enhance regions with high pixel variability.

A manual threshold was applied to refine the segmentation: pixels with intensity values between 10 and 255 were considered as foreground, while those below 10 were designated as background. Finally, the images were converted to binary masks. The background was set to black to facilitate clear segmentation of the grid and objects in the binary mask.

The BF images were stitched with the MIST plugin [5], [6]. Images were divided in two sets, each stitched separately. The stitched images were cropped to a uniform width and length to ensure compatibility in the final stitching. The two cropped images were then stitched together to create a single composite image. This three-step stitching allowed for better image assemblies than assembling all the images at once (based on visual observation). At each stitching, a text file containing the relative coordinates of images was generated and saved for later assembly of BF and Green Fluorescent (GF) mega pictures.

3. 3. Stitching review

The transformed BF images of the 10 fields were stitched using predetermined coordinates defined *via* the MIST plugin with pre-stitching for optimal alignment as described above. Cropping of the megapictures was performed to center images, normalize pixel count, and remove edge artifacts.

A visual inspection of all stitched images was performed to ensure the accuracy of the mega picture reconstruction. Some instances of redundancy or excessive cropping between images were identified, but these affected only a small number of pictures and of pixels. These minor stitching defects were independent of the presence of plastic particles and also independent of the GFP signal, as the reconstruction was based solely on BF pictures.

After cropping, the green fluorescence images were organized into stacks to replicate the depth of field.

3. 4. Segmentation

Segmentation of MPs employed trained models *via* Fiji's trainable Weka segmentation plugin [7], [8], a machine learning-based image analysis. Given the shape heterogeneity of particles and their refractive properties, this plugin enabled a fine and reproducible segmentation of microparticles. The segmentation resulted in binary pictures capturing the position of the MPs on the stitched grayscale-transformed BF picture.

The GF pictures were split into the three color channels, and only the green channel was retained for further analyses. The segmentation of fluorescent bacteria was performed differently for various polymers. For PS and glass beads, a model was trained using the Trainable Weka Segmentation module to differentiate pixels associated with fluorescent bacteria from those associated with light passage through the center of beads.

For PP samples, bacteria were segmented by applying a threshold of fluorescence intensity of 82. For PET samples, bacteria attached to microplastic particles (MP) were segmented by applying a threshold of 90, converting the images to binary with a dark background, and saving the masks. Planktonic bacteria were segmented by subtracting areas of MP using a BF mask, setting an automatic threshold, then applying a threshold of 82, converting to binary, and saving the masks. We used a higher segmentation threshold for bacteria overlapping with MP because of autofluorescence of PET particles, resulting in a high subsequent proportion of falsely segmented pixels.

Each mask produced by its utilization was visually verified by overlaying it onto the original image. Manual corrections were made in cases of local mis-segmentation. Cases of mis-segmentation for the brightfield mask included pixels falsely classified within the mask (Malassez cell grid, slide background) or falsely classified outside the mask (transparent areas of MP).

Given the heterogeneity of particles in shape and refractive properties, this plugin enabled a fine and reproducible segmentation of microparticles. Image processing had to be fine-tuned to each polymer to account for differences of properties between them. For fluorescence images, it was necessary to increase the threshold for PET samples due to its autofluorescence, and to apply a trained model for PS and glass beads, which exhibited fluorescent centers at certain depths.

3. 5. Glass beads

For glass bead images, a bandpass filter was applied on 8-bit grayscale pictures with parameters set to filter out structures larger than 100 pixels and smaller than 20 pixels, with a tolerance of 5. Models were trained using the Trainable Weka Segmentation module to obtain a microparticle mask, and differentiate pixels associated with fluorescent bacteria from those associated with light passage through the center of the beads (supplementary material). Fluorescent pixels overlapping those of the MPs and present from 200 to 100 μm were classified as originating from bacteria attached to the glass beads.

##### 3.6. Fiji pipeline evaluation

For each polymer, negative fluorescence controls were conducted to verify that each analysis pipeline did not contribute to any additional fluorescence signal, thereby ensuring that the detected fluorescence was exclusively attributable to the fluorescent bacteria. Briefly, sterile microplastics were placed on a Malassez cell, and images of the fields were captured as described in the Materials and Methods section. The dedicated pipeline was then applied to these controls, following the rules and parameters corresponding to the relevant polymer. Each condition was done in four replicates.

The average sum of fluorescent pixels detected over the ten layers for each sample is given in the table below (± standard deviation):

| Polymer | Fluorescent pixels overlapping MPs | Fluorescent pixels not overlapping MPs | Not fluorescent pixels forming MPs |
| --- | --- | --- | --- |
| PS | 31 (± 34) | 12 (± 14) | 5,982,756 (± 1,773,946) |
| PP | 741 (± 211) | 1,668 (± 717) | 4,419,720 (± 1,137,888) |
| PET | 908 (± 295) | 1,392 (± 624) | 5,265,518 (± 1,743,584) |
| GL | 287 (± 118) | 26 (± 29) | 5,225,810 (± 1,314,943) |

We performed a linear regression analysis to investigate whether there was a significant difference in the total number of fluorescent pixels detected across different polymers, taking into account the two types of fluorescent pixels (planktonic and attached). No significant effect was observed, even when including the interaction between polymer type and fluorescent pixel type.

To assess the existence of a pipeline-specific bias in the segmentation of fluorescent bacteria between polymers, images of fluorescent bacteria without microparticles were reused from five samples. The estimated number of bacteria in each sample was ~100. These pictures were processed using the three types of fluorescent pixel segmentation developed to account for the specific characteristics of each polymer. The average sum of fluorescent pixels detected over the ten layers for each sample is given in the table below (± standard deviation):

| Fluorescent segmentation analysis | Number of pixels classified as fluorescent |
| --- | --- |
| PS (Trainable Weka model) | 1,700 (± 949) |
| PP/PET (setting intensity threshold to 82) | 2,579 (± 848) |
| GL (Trainable Weka model) | 2,866 (± 1,717) |

We performed a linear regression analysis to investigate whether there was a significant difference across the segmentation analyses. No significant effect was observed (linear regression analysis, p-values > 0.14), meaning that all pipelines process fluorescent bacteria consistently, ensuring that the signal from bacteria is directly comparable.

### 4.  Experiment 2: *V. anguillarum* attachment on field-colonized microplastics

###### Sampling device

We prepared the sampling device as described in [9]. For each incubation, three nylon bags (10 cm x 6.5 cm, 25 µm mesh) containing PP, PS, or PET microparticles (50 mg) were glue-sealed and hung with Ty-raps within an oyster cage (11.5 x 25.5 x 72 cm, Nodus factory, France). The bags were infused in sterile water in the lab to remove all microplastics < 25 μm diameter through the mesh and check for leaks, thus preventing any plastic contamination of field sites. The cage was loaded with four 1 kg diving weights and immersed one meter below the surface. The incubation period was four weeks.

###### Field-colonized microplastic collection

After incubation, the bags containing MPs were collected and processed in the lab as described in [9]. The external side of the bags was scraped to remove potential contamination. The closed bags were subsequently bathed in 50 mL of sterile artificial seawater (Sea salt, InstantOcean, VA, USA) to remove particles and microorganisms < 25 µm. The bags were then cut open and the content was resuspended in a sterile solution by aspersion. The obtained solutions were transferred onto a 20 µm mesh filter to collect MPs and discard planktonic material. The field-colonized MPs were resuspended in 2 mL of artificial seawater, of which 400 µl were dedicated to *V. anguillarum* attachment rate measurements.

### 5.  Statistical analyses

Model parameters were estimated in a Bayesian framework, using a Markov Chain Monte Carlo (MCMC) algorithm, implemented with the R package brms [10], [11], [12]. For each model variant, 4 chains were run for 20,000 iterations each. 8,000 values were discarded as burn-in, such that convergence of the Markov chains was reached. We sampled every 50^th^ value as this was enough to avoid autocorrelation within the Markov chains. A Gelman-Rubin statistic below 1.1 for all parameters was considered to indicate that between-chain variance was low enough compared to the within-chain variance. Non informative priors were used for all parameters. The models were compared using the Watanabe-Akaike Information Criterion (WAIC). Among the models with the smallest WAIC values and a difference of less than 4 units, we selected the most parsimonious model.

Supplementary Results

### 1. Phylogenetic analyses

Strains were isolated in three main areas: the northern European marine area (n= 8), the Mediterranean Sea (n= 6), and fish farming in the United States (n= 2) (Figure 1A). The number of SNPs between strain pairs isolated from the Mediterranean Sea, as well as between the two strains from the United States, was lower than that observed between strains originating from different geographic regions (39,505 SNP, 484 SNP, and 50,721 SNP, respectively). Conversely, strain pairs from Northern Europe exhibited a higher number of SNPs than inter-regional comparisons (51,739 SNP), suggesting that geographic proximity does not consistently reflect phylogenetic relatedness.

### 2. *V. anguillarum* attachment on pristine microplastics (experiment 1)

Four samples missed one brightfield image each, and one sample missed five brightfield images due to technical issues. Consequently, these samples had a reduced number of processed microscopic fields, leading to a decrease in the number of pixels analyzed, ranging from 7.5% to 52% relative to the average pixel count. For six samples, one or two green fluorescence images were missing in the two least deep layers (160 µm and 180 µm above the slide). After ensuring that no fluorescent pixel was detected on the remaining pictures from the same layers, we replaced the missing image one by copying another one from the same layer, before applying our Fiji pipeline.

Two of the affected samples were strains incubated with PS, five with PP, and three with PET (for different strains each time, Supplementary Table S4). Despite these issues, the samples were retained in the analysis, and no sample was excluded.

There was no significant difference in picture sizes between polymers (anova, p-value= 0.06, Figure S2).

We noted a sharp decrease in the number of fluorescent pixels detected on PS pictures compared to PET and PP (PET= 96,014 fluorescent pixels/sample ± 233,769, PP= 15,244 fluorescent pixels/sample ± 16,468, PS= 3,261 fluorescent pixels/sample ± 3,073). This was confirmed by visual inspection *via* microscopy, where we saw only minor bacterial attachment to PS microparticles compared to the other polymers.

### 3. *V. anguillarum* attachment on field-colonized microplastics (experiment 2)

One sample missed one microscopic field. Therefore, the number of pixels analyzed was 25% less than the average pixel count. Two samples had two columns of five fields merged into a megapicture, due to the lack of microparticles on the slide. However, the number of pixels analyzed was similar to that of samples processed as described in the Materials and Methods section. For four samples, one or two green fluorescence images were missing in the two least deep layers (160 µm and 180 µm above the slide). After ensuring that no fluorescent pixel was detected on the remaining pictures from the same layers, we replaced the missing image by copying another one from the same layer, before applying our Fiji pipeline.

Two of the affected samples were strains incubated with PS, two with PP, one with PET, and two with GL (Supplementary Table S4). Despite these issues, the samples were retained in the analysis, and no sample was excluded.
