## Supplementary figures and images for "The attachment rate of *Vibrio anguillarum* strains to microplastics strongly varies with abiotic and biotic factors"

### Supplemental Figure 1

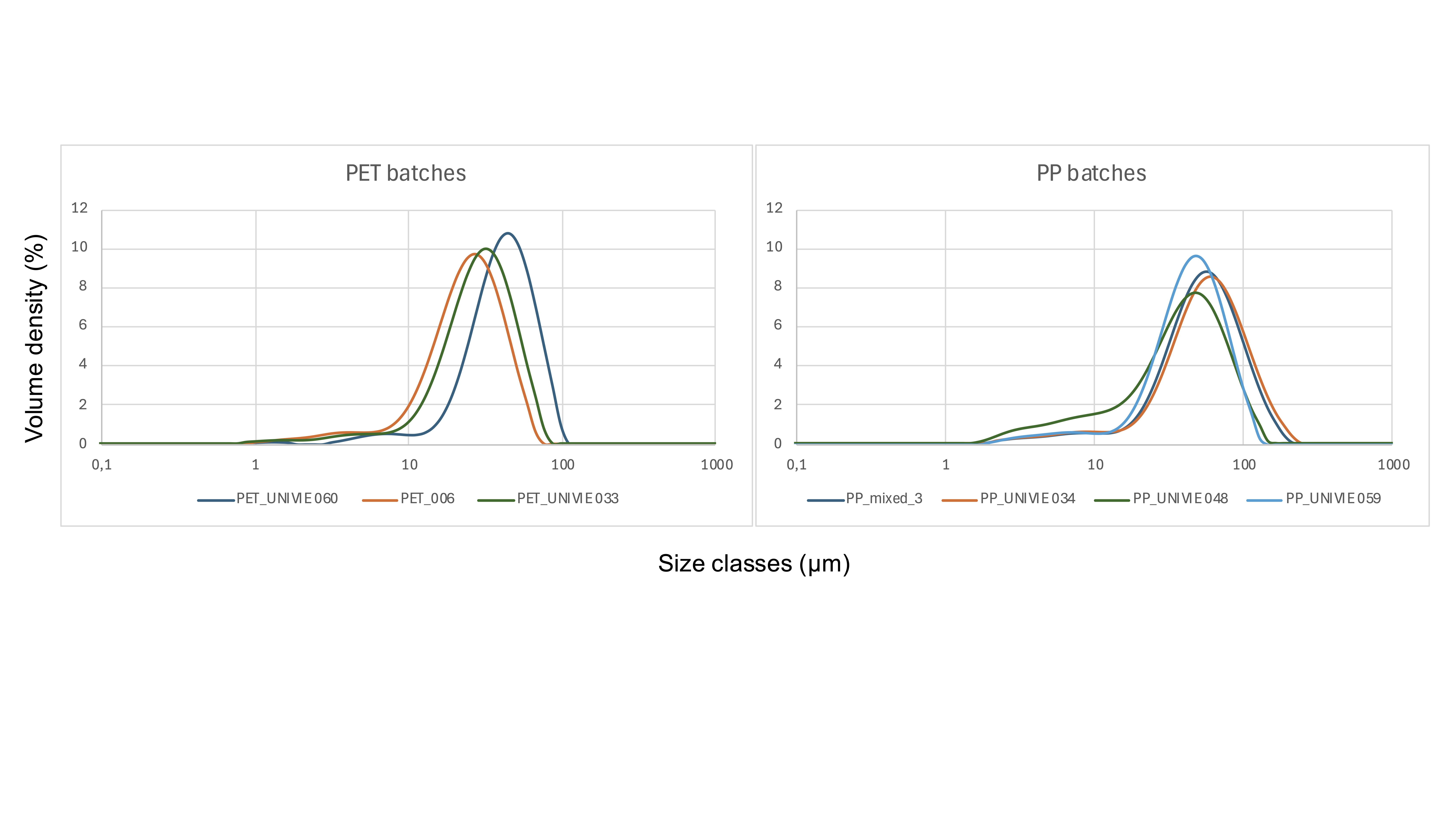

### Supplemental Figure 2

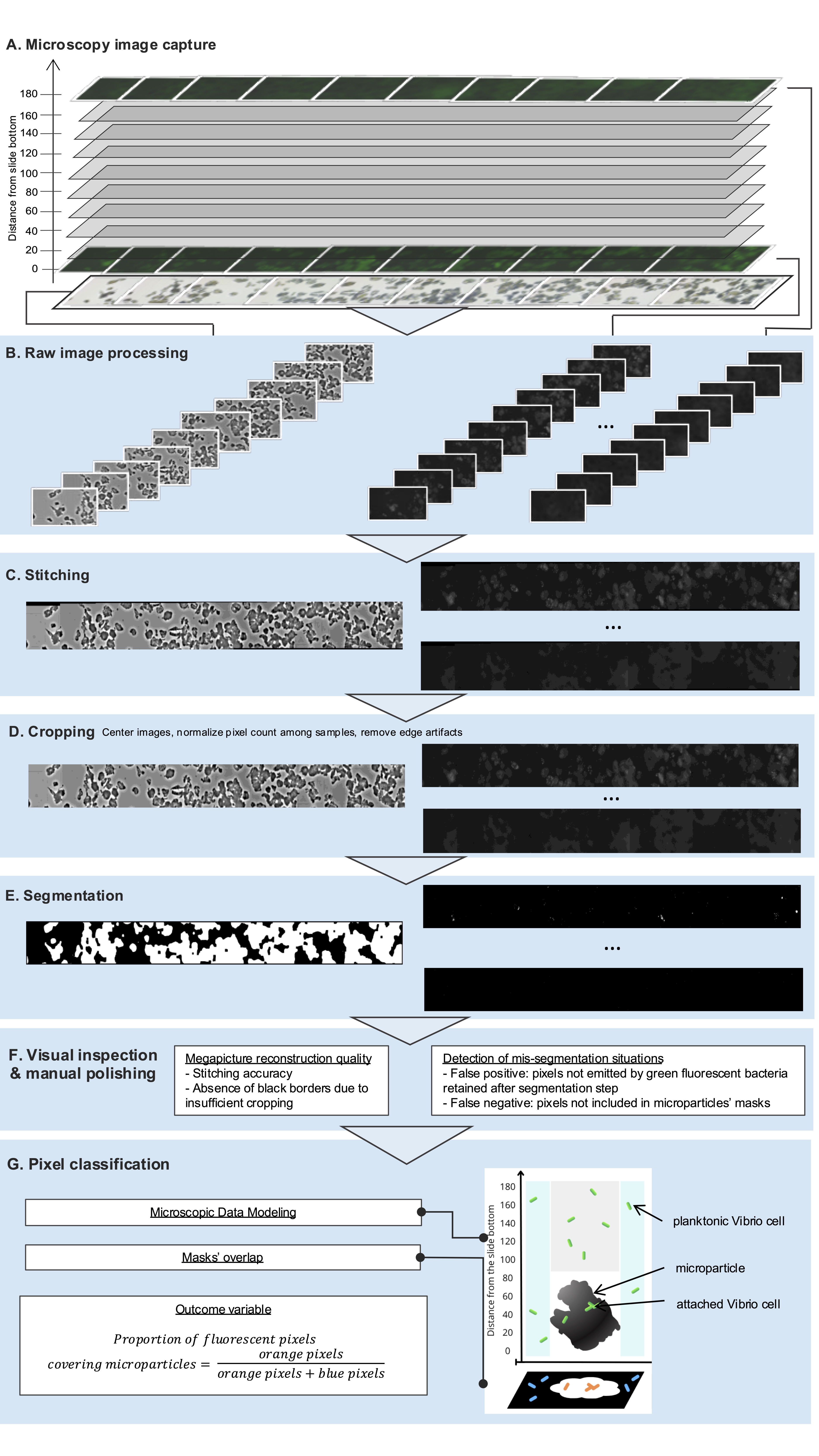

### Supplemental Figure 4

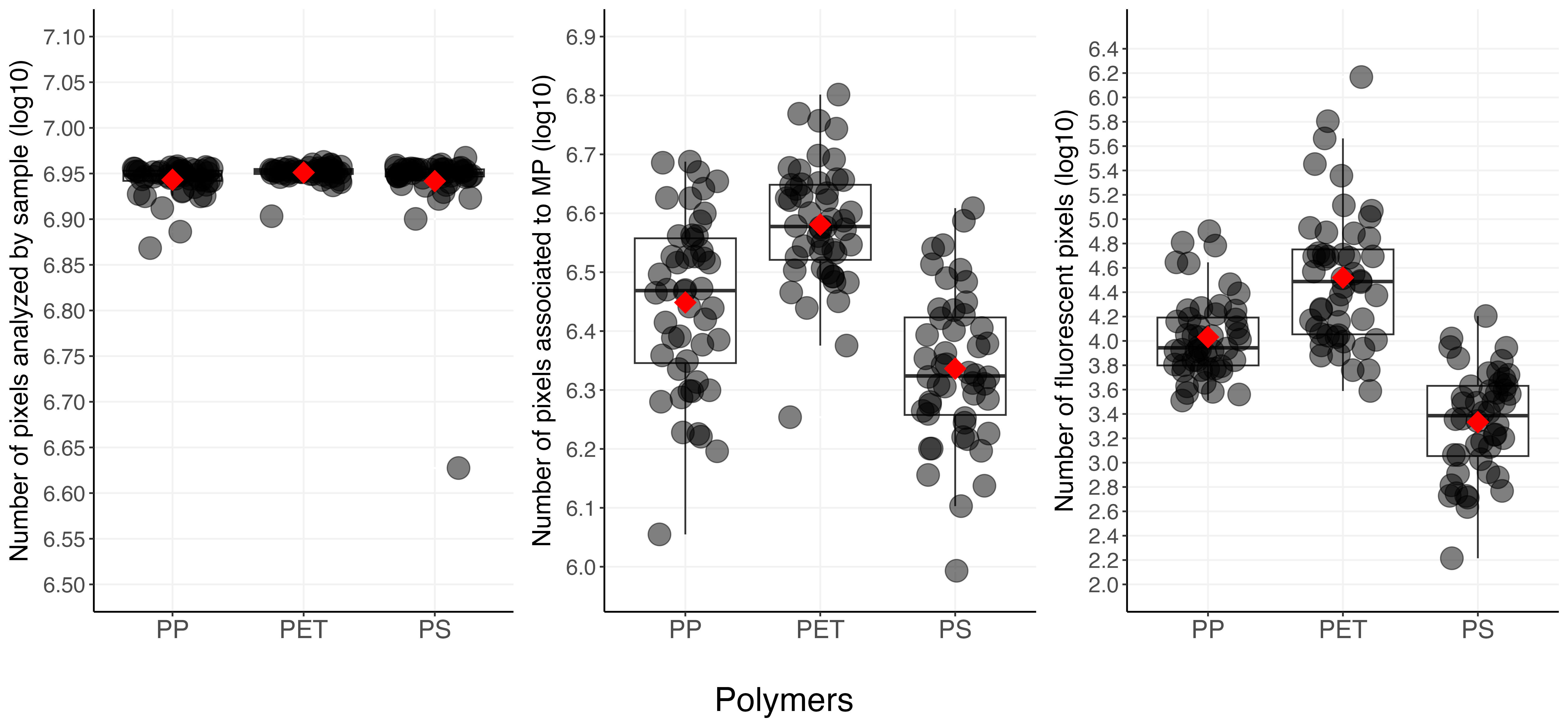

### Supplemental Figure 5

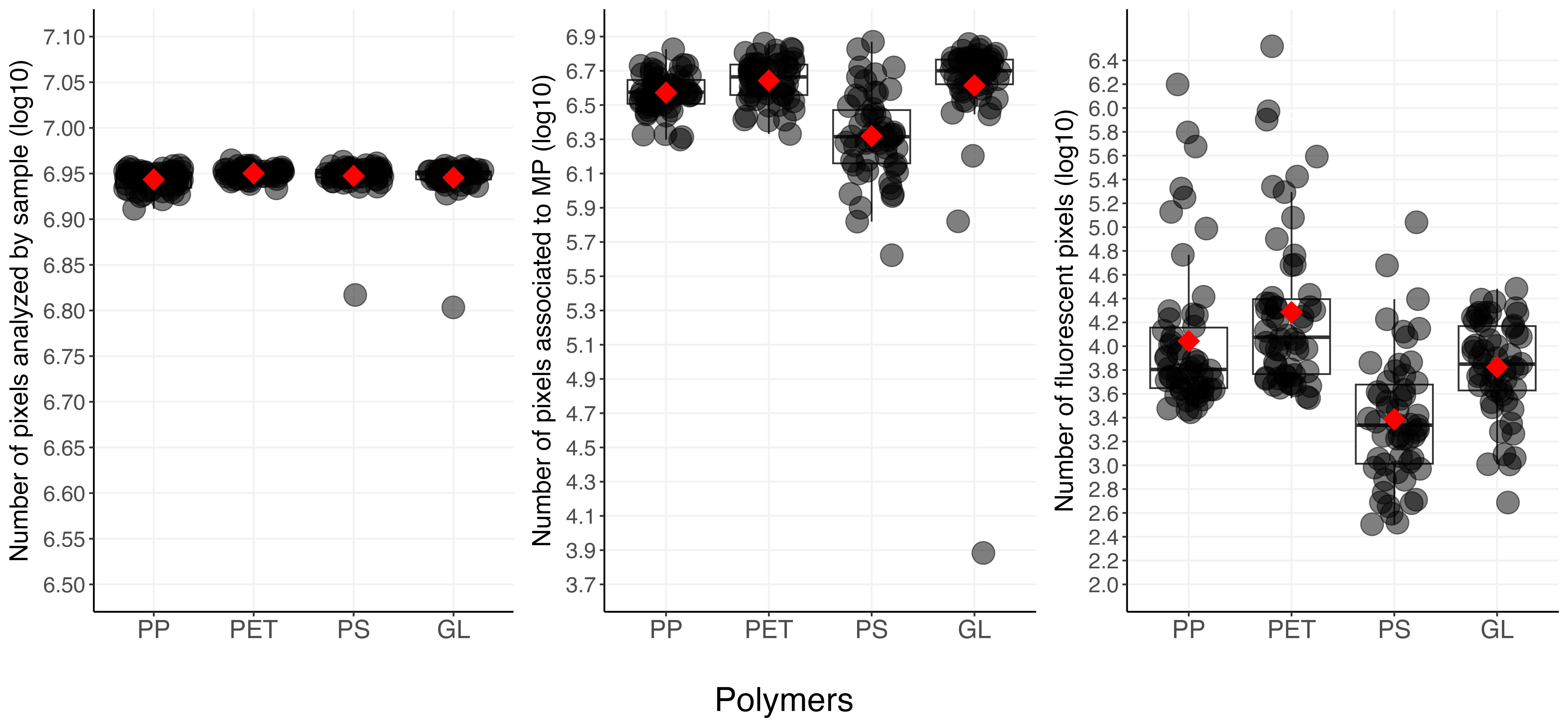

### Supplemental Figure 6

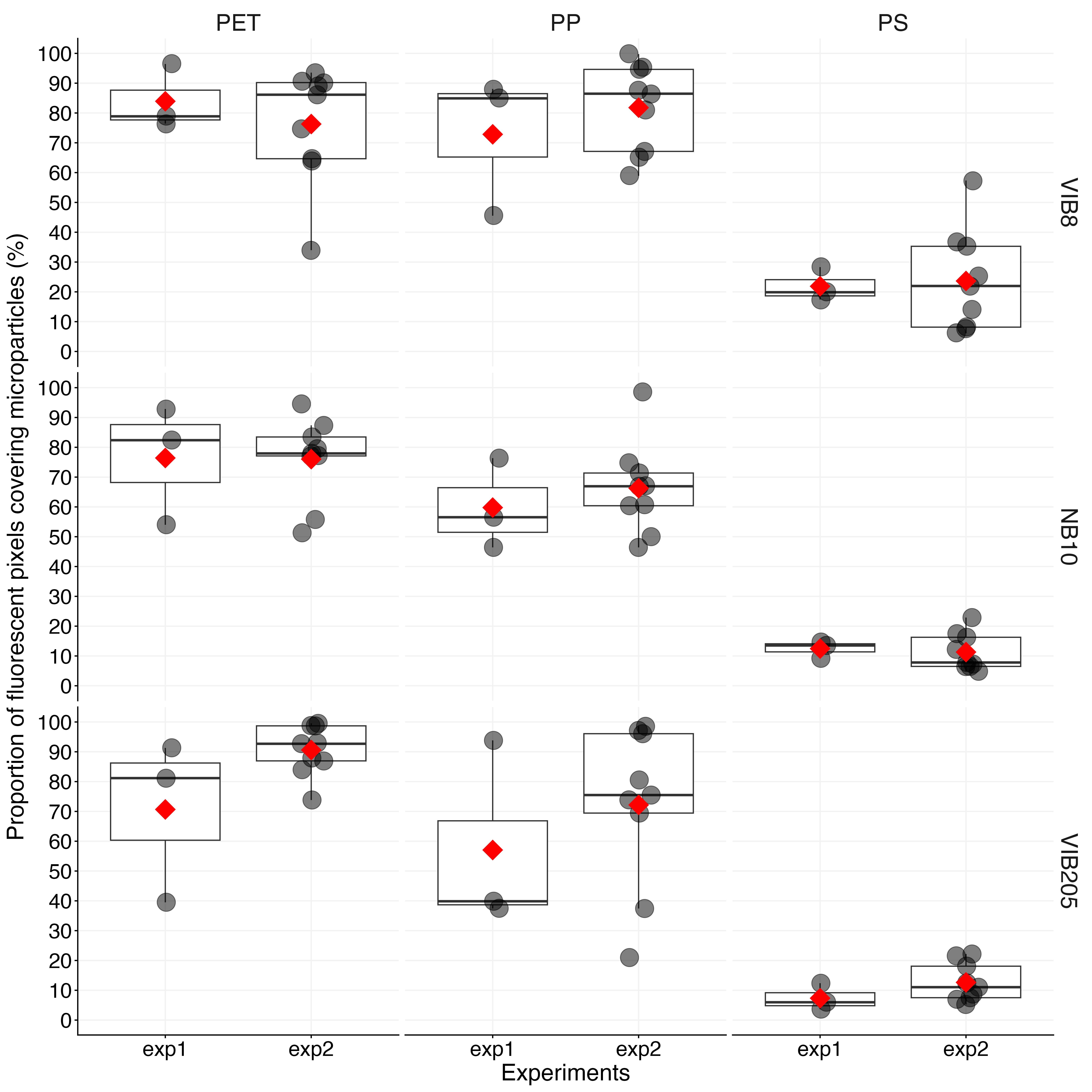

### Supplemental Figure 7

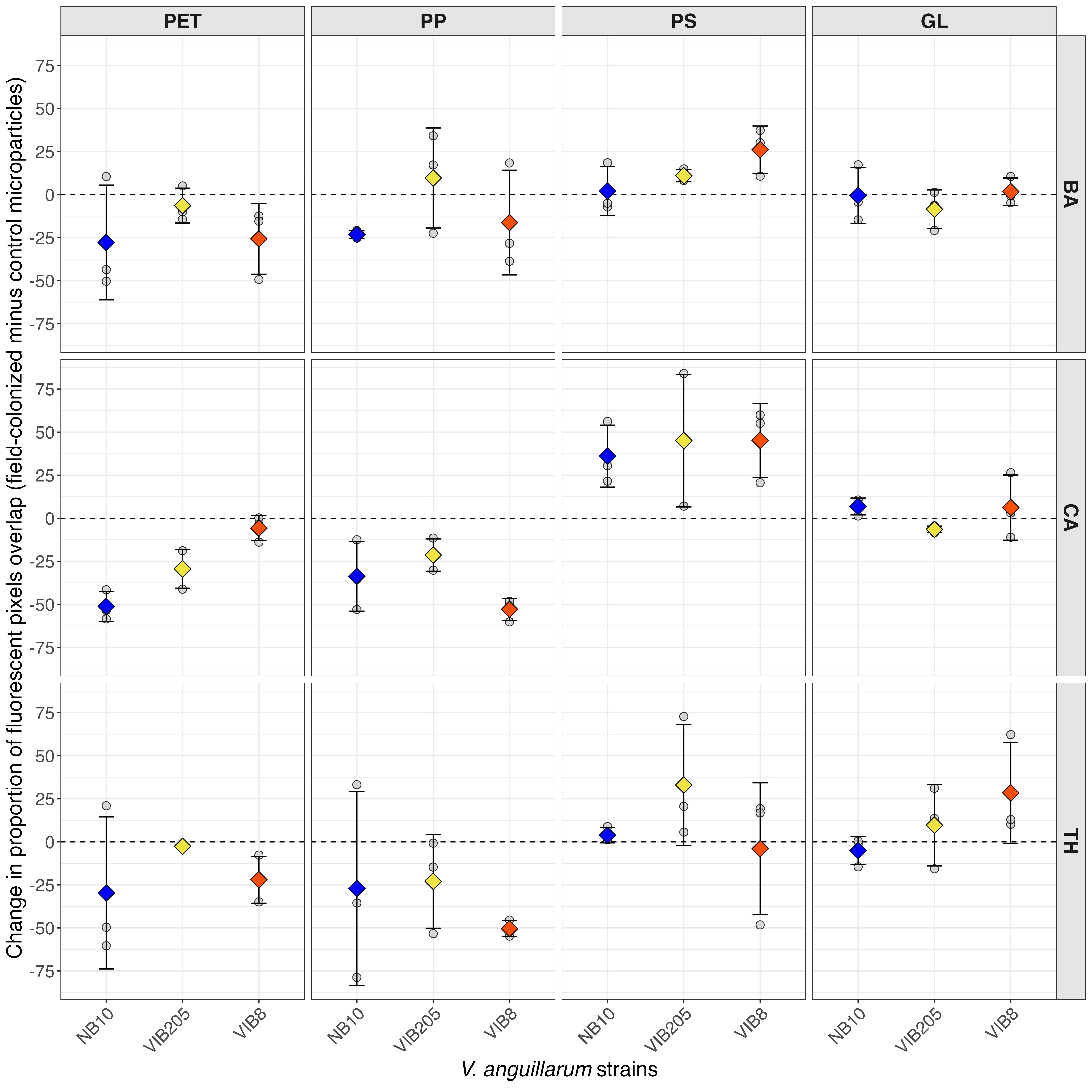
